# Towards *In Situ* Dynamics of DNA-bound Full-Length p53 Tetramer

**DOI:** 10.1101/2025.10.14.682466

**Authors:** Özlem Demir, Emilia P. Barros, Rommie E. Amaro

## Abstract

p53 is the most important tumor suppressor in humans as well as the most frequently mutated gene found in human cancers with ∼50% of all human tumors bearing p53 missense mutations that leave p53 inactive. Restoring the p53 activity proved to lead to tumor regression even in advanced tumors in mouse models— and thus, is among the most attractive potential strategies for novel cancer therapy. Full-length p53 (fl-p53) consists of 393 residues and multiple domains; some folded and some disordered. Using crystal structures of folded domains and integrative molecular modelling techniques for disordered domains, we generated the first wild-type fl-p53 tetramer model bound to DNA. When solvated, the system size nears 500K atoms challenging extensive sampling. Using Anton2 supercomputer for microsecond-timescale simulations in explicit solvent and the rigorous Markov state model (MSM) framework, we elucidated the conformational landscape of wild-type p53 as well as two of the p53 hot-spot cancer mutants, Y220C and G245S, in a physiological DNA-bound, full-length tetramer context. In the simulated timescale, DNA-bound fl-p53 tetramer bent DNA and formed a compact complex with interactions between the N-terminal and DNA-binding domains (DBDs), and the C-terminal domains (CTDs) with DNA. WT fl-p53 tetramer also sampled a unique quaternary DBD organization not accessed by the cancer mutants. Free energy landscapes indicated differential dynamics for inner and outer p53 DBDs due to the dimer-dimer interface. The dynamics of the druggable L1/S3 pocket is also closely monitored. Ultimately the MSMs identified an underexplored loop 6 (L6) cryptic pocket and captured the effect of p53 tetramerization and cancer mutations.

**Significance:** p53, the most important tumor suppressor in humans, is found inactivated due to single point mutations in almost 50% of all human tumors. As restoring p53 activity is shown to achieve tumor regression even in advanced tumors in mice, there is a lot of interest in finding small molecules to reactivate p53 cancer mutants. p53 has 393 residues and is an intrinsically disordered protein with multiple unstructured and dynamic domains that are known to interact with and help the function of its folded domains. Adding another level of complexity, p53 forms a tetramer to bind DNA and start its transcriptional activity. Due to its high dynamicity, p53 evades full structural characterization leaving many open questions. We generated an integrative model weaving together structures of isolated p53 domains and explored it with molecular dynamics simulations at microsecond timescale. This allowed us to monitor various unresolved aspects with atomistic detail like the dynamics of druggable pockets, the interplay of p53 domains in the physiological complex and how they change in p53 cancer mutants compared to the wild-type p53. Our study provides insights into the structure and dynamics of a key tumor suppressor protein at its physiologically relevant state of full-length tetramer bound to DNA.

## INTRODUCTION

p53 is arguably the most important tumor suppressor in humans, and also the most frequently mutated gene found in human cancers, with ∼50% of all human tumors bearing p53 missense mutations that leave p53 inactive^1^. Almost all these mutations are localized at the p53 DNA-binding domain (DBD), emphasizing the vital role of DBD. A large spectrum of cellular and tissue activities is known to be controlled by p53, such as induction of cell cycle arrest, apoptosis, senescence, autophagy, angiogenesis, cell migration, suppression of cancer cell specific metabolism, and promotion of anti-tumor microenvironments, which collectively prevent tumor initiation and maintenance^2^. In addition to its transcription factor role by which p53 controls expression of many protein-coding genes as well as microRNAs, recent findings highlighted the importance of transcription-independent functions of p53 in tumor suppression as well^3^. Regardless of the mechanism, restoring the p53 activity proved to lead to tumor regression even in advanced tumors in mouse models^4–6^— and thus, is among the most attractive strategies for novel cancer therapy^7^. The promising clinical trial results of Y220C cancer mutant reactivator molecules^8^, which were based on small molecules identified in the Fersht lab by targeting a mutant-specific druggable pocket^9^, raise hopes for such a strategy.

The structural characterization of full-length intrinsically disordered proteins (IDPs) such as p53 that consist of both folded and disordered domains is a major challenge. While performing cellular functions, folded or disordered components of multi-domain IDPs do not function as isolated entities, but instead all components of the entire protein act in synergy. p53 consists of 393 residues that are classified into an N-terminal transactivation domain (NTD), a DNA binding core domain (DBD), a linker region, a tetramerization domain (TET) and a C-terminal domain (CTD), respectively (Fig 1A). Among these, only DBD and TET are folded while the rest of the protein is mainly disordered. P53 binds DNA as a tetramer (dimer of dimers). Even though there are many crystal structures available for various isolated domains of full-length (fl-p53), there is still no x-ray crystal or NMR structure of fl-p53 tetramer in complex with DNA. p53 DBD tetramer in complex with DNA is the largest region that has been resolved^10,11^. Using crystal structures of folded domains and integrative molecular modelling techniques for disordered domains, we previously generated the first wild-type (WT) fl-p53 tetramer model bound to DNA (Fig 1B) and explored its dynamics with triplicate ∼100 nanosecond MD simulations for three different DNA sequences^12^. Characterizing the structural ensembles and dynamics of the fl-p53 tetramer while in complex with DNA in atomic detail is invaluable for understanding how p53 operates, as well as understanding how they change in the case of a p53 cancer mutation.

**Figure 1.**
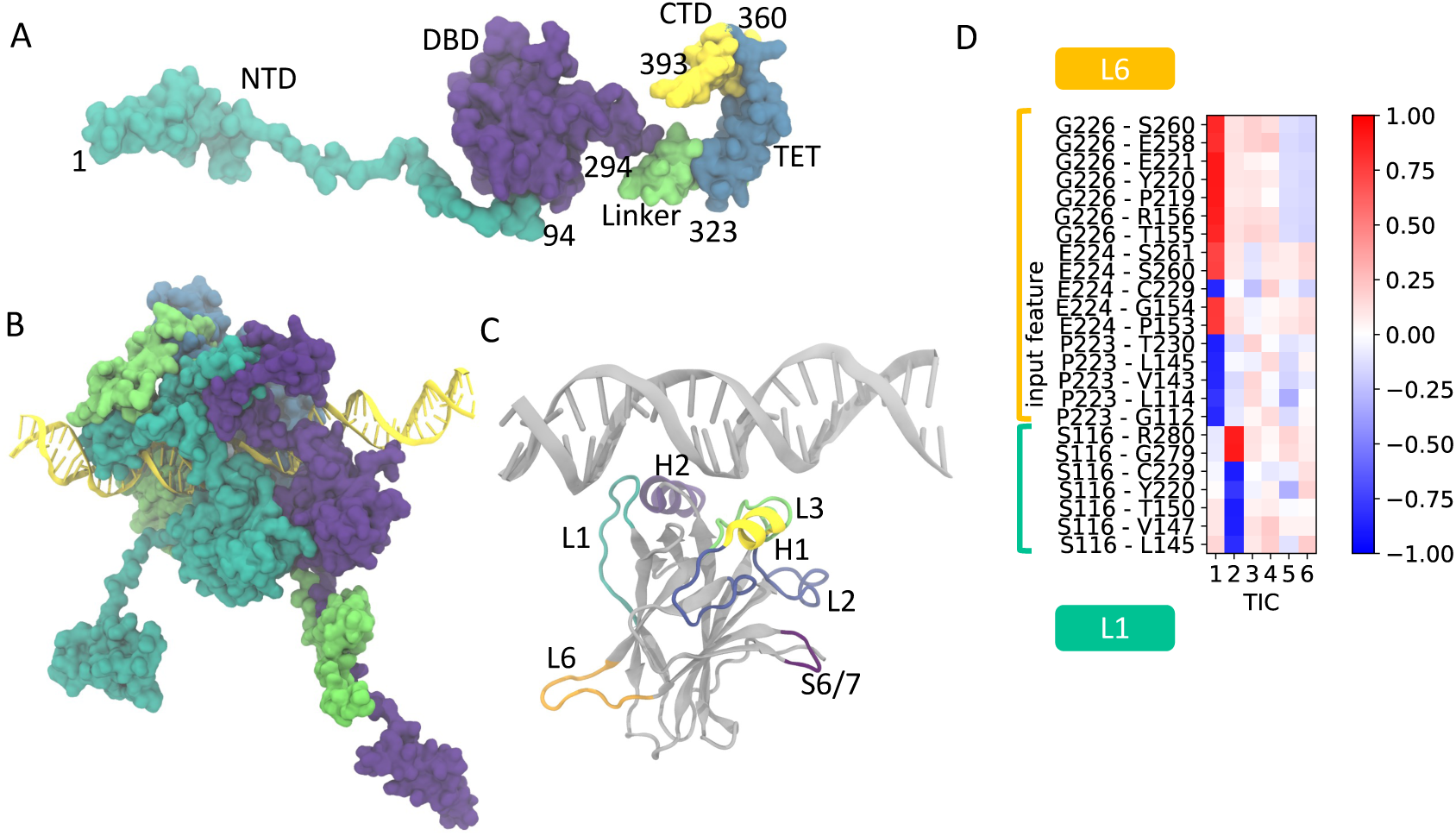
A) Different domains of full-length p53 monomer. Starting structure for monomer A is depicted. B) DNA-bound full-length p53 tetramer structure. Starting structure for MD is depicted. C) DNA-bound p53 DBD important loops and helices. Chains B, E and F from pdbID:1TSR are depicted. D) Correlation of features with the first six tICA components. Input features, which are pairwise distances that involve L1 or L6 anchor residues, are depicted. The transitions involving L6 loop are slowest while those involving L1 are the second slowest.

In this study, we extended the explicit-solvent MD simulations of WT fl-p53 tetramer bound to the p21 response element DNA on the microsecond timescale. We additionally ran comparable MD simulations for the Y220C and G245S mutants. To our knowledge, these are the longest timescale simulations to date for these systems in full-atom detail. Using Anton2^13^ supercomputer for long-timescale simulations and the state-of-the-art MD/MSM analysis framework, we elucidated the conformational landscape of p53 for WT and two of the hot-spot cancer mutants, Y220C and G245S, in a DNA-bound full-length tetramer context. We paid close attention to the druggable L1/S3 pocket, which is a cryptic druggable pocket that is universally shared across all p53 mutants.

## RESULTS

We performed explicit-solvent molecular dynamics (MD) simulations of the DNA-bound fl-p53 tetramer. For starting coordinates, we used the final frame of 100-ns MD simulation of the DNA-bound WT fl-p53 tetramer model in our Oncogene 2017 study^12^. Three systems are simulated in this study: wild-type, Y220C and G245S p53 mutants. Each system was simulated for 3 independent replicas of 7.5 µs, with coordinates recorded every 0.24 ns, adding up to a record sampling of 270 µs (7.5 µs x 3 replicas x 4 monomers x 3 systems).

Root-mean-square-deviation (RMSD) analysis of the backbone atoms of all four p53 DBDs of the tetramer with respect to their initial coordinates indicate stability of the simulations (Fig S1). Secondary structure analysis of p53 DBD proves the overall protein fold is conserved throughout simulations (Fig S2).

In the root-mean-square-fluctuation (RMSF) plots of the MD trajectories (Fig S3), the NTD is the most flexible and the DBD is the most stable domains of fl-p53 for all systems. With a closer look at the DBD (Fig S4), the most flexible regions captured in RMSF analysis are L1 loop, S6/S7 loop and L6 loop across all monomers in all three systems. Both L1 and L6 loops in p53 are part of transiently open cryptic druggable sites^9,14–20^.

### DNA, CTD and NTD Dynamics in fl-p53 Tetramer Simulations

Experimentally, fl-p53 tetramer was measured to bend the DNA it binds to by 27-40° ^11,21–23^. In line with experimental results, DNA bending angle in fl-p53 tetramer simulations is calculated to be 37 ± 9°, 45 ± 9° and 28 ± 11° (ensemble average ± standard deviation) for the WT, Y220C and G245S systems, respectively (Fig S5).

An interesting observation in the simulations is the significant change towards a more compact form of fl-p53 tetramer, which is reflected in the marked reduction of the radius of gyration of the fl-p53 tetramer in all MD copies from the initial 50 Å to as low as 38 Å in multiple replicas (Fig 2, Fig S6). This change is partly due to the interaction of the lysine-rich CTD with DNA outside the p53 recognition element. Using ^15^N-labeled CTD domain in NMR of fl-p53, the CTD domain was shown to loosely tether p53 to the DNA through weak, sequence-independent and highly dynamic contacts (involving residues 369−374 and 381−388, especially K372, K373, and K382) with the phosphate backbone of DNA^24^. In our MD simulations of all 3 systems, we observe highly dynamic DNA interactions of positively charged residues in p53 CTD including K372, K373 and K382 as plotted in Fig S7.

**Figure 2.**
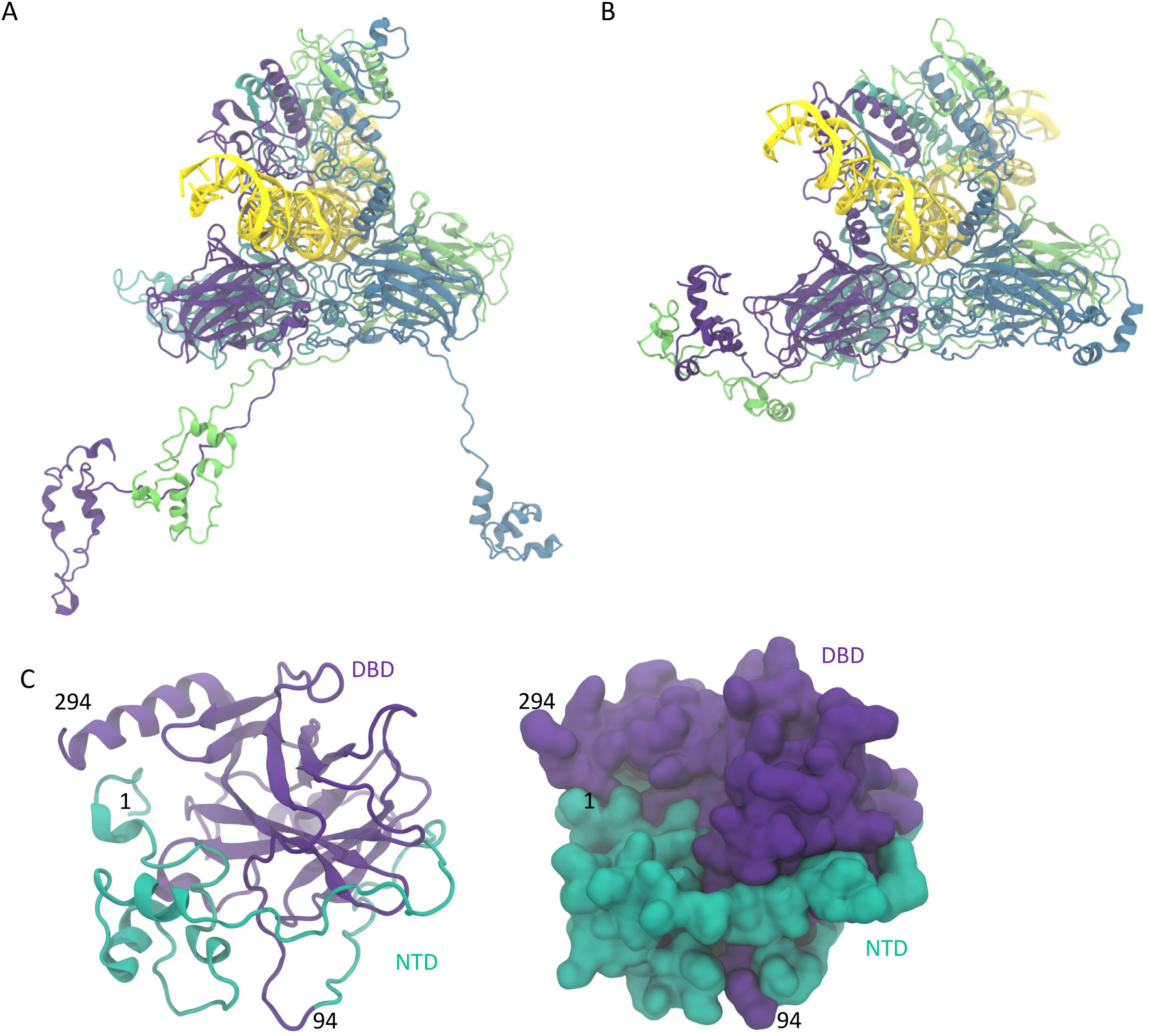
WT fl-p53 tetramer conformational change observed in MD simulations. A) Conformation with lowest number of NTD-DBD contacts. B) Conformation with highest number of NTD-DBD contacts. DNA and p53 tetramer are depicted in ribbons, with each p53 monomer colored differently in panels A and B. C) Snapshot of WT monomer B conformation with highest number of NTD-DBD contacts observed in MD simulations. DBD in purple, NTD in green. Only DBD and NTD domains are depicted for simplicity (in ribbons as well as in surfaces).

However, the overall compactness of fl-p53 we observed in the MD simulations is mostly due to the dynamic and largely hydrophobic interactions of the NTD residues with DBD residues (Table S1). DBD and NTD that belong to the same monomer interact with each other in all MD replicas while less pronounced DBD-NTD interactions between different monomers are also observed (Figure S8, S9, S10). The plots also indicate more extensive NTD-DBD interactions in the inner monomers B and C in all systems. Quantitative analysis of the interactions shows that the residues that interact with the DBD lie across the NTD (Fig S11). Our findings are in line with single-molecule FRET studies of fl-p53 tetramer from the Fersht lab^25^ as well as the NMR studies of He et al which showed that the NTD domain (residues 40-61 and 64-92) of p53 interacts with the DBD domain^26^. While this manuscript was in preparation, the first full-length cryo-EM structures of the fl-p53 monomer and dimer in the absence of DNA with 4-5 Å resolution (Pdb ID 8F2I and 8F2H) were published, further proving direct DBD-NTD interactions^27^.

### p53 Organization around DNA

Next, we explored if the WT and mutant p53 tetramers differ in their quaternary binding modes to DNA in microsecond timescale MD simulations. A combined principal component analysis (PCA) of the C-alphas of fl-p53 tetramer DBD finds a significant difference between the three systems (Fig 3). First, G245S mutant samples high PC1 values not accessed by the other two systems (Fig 3). PC1 corresponds to the motion between symmetric tetramer DBDs (low PC1 values) to asymmetric tetramer DBDs (high PC1 values) (Fig 4A). Comparison of the structures at each extreme of the PC1 scale highlight that, from lowest to highest PC1 values, the following conformational changes occur: 1. both monomer B and monomer D DBDs move further away from DNA, 2. monomer A and monomer D DBDs move away from each other, 3. monomer B and monomer C DBDs approach each other (Fig 4A). The structure at high PC1 value displays a loose/non-optimal DNA binding mode similar to what we observed previously for WT p53 nonspecific-DNA binding as well as for DNA response element binding of p53 cancer mutants^28,29^. Interestingly, Ser183 in monomer C and Asp186 in monomer B interact directly in the structures that correspond to highest PC1 value while these two residues are 17 Å apart in the structures that correspond to lowest PC1 value in the G245S system (Fig. S12 A). Direct interaction between monomer B Asp186 and monomer C Ser183 is observed continuously for about 5 μs in MD replica 2 while visited briefly at different times in the other two G245S MD replicas (Fig. S12 B).

**Figure 3.**
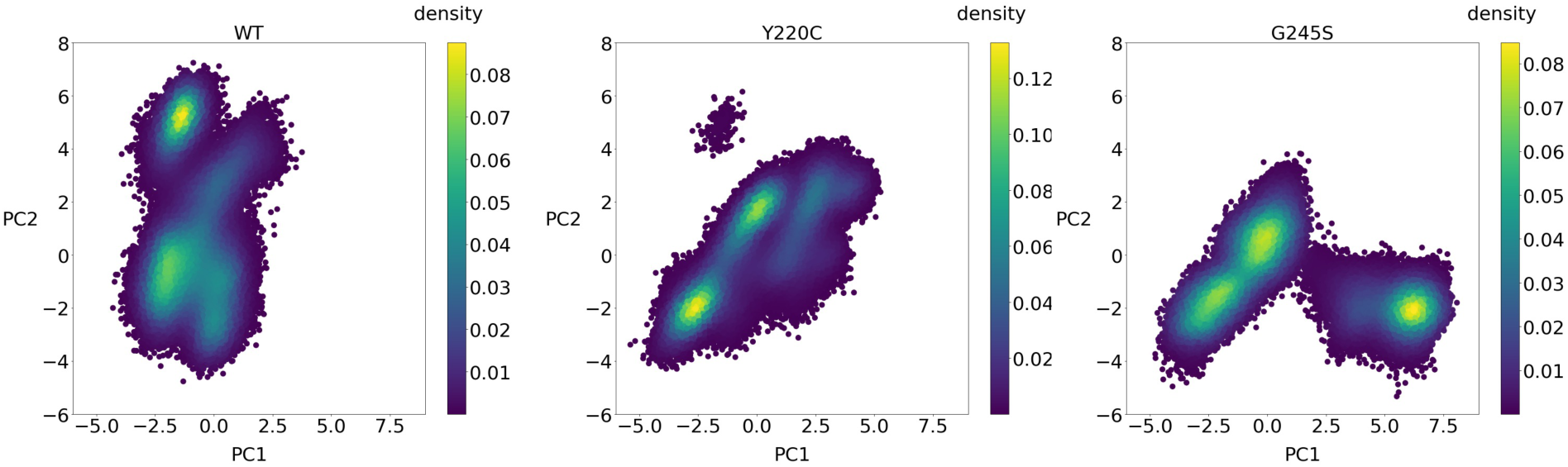
Principal component analysis (PCA) of fl-p53 DBD tetramer C-alphas. The MD starting structure projects to PC1=-0.41, PC2=0.27 in this space.

**Figure 4.**
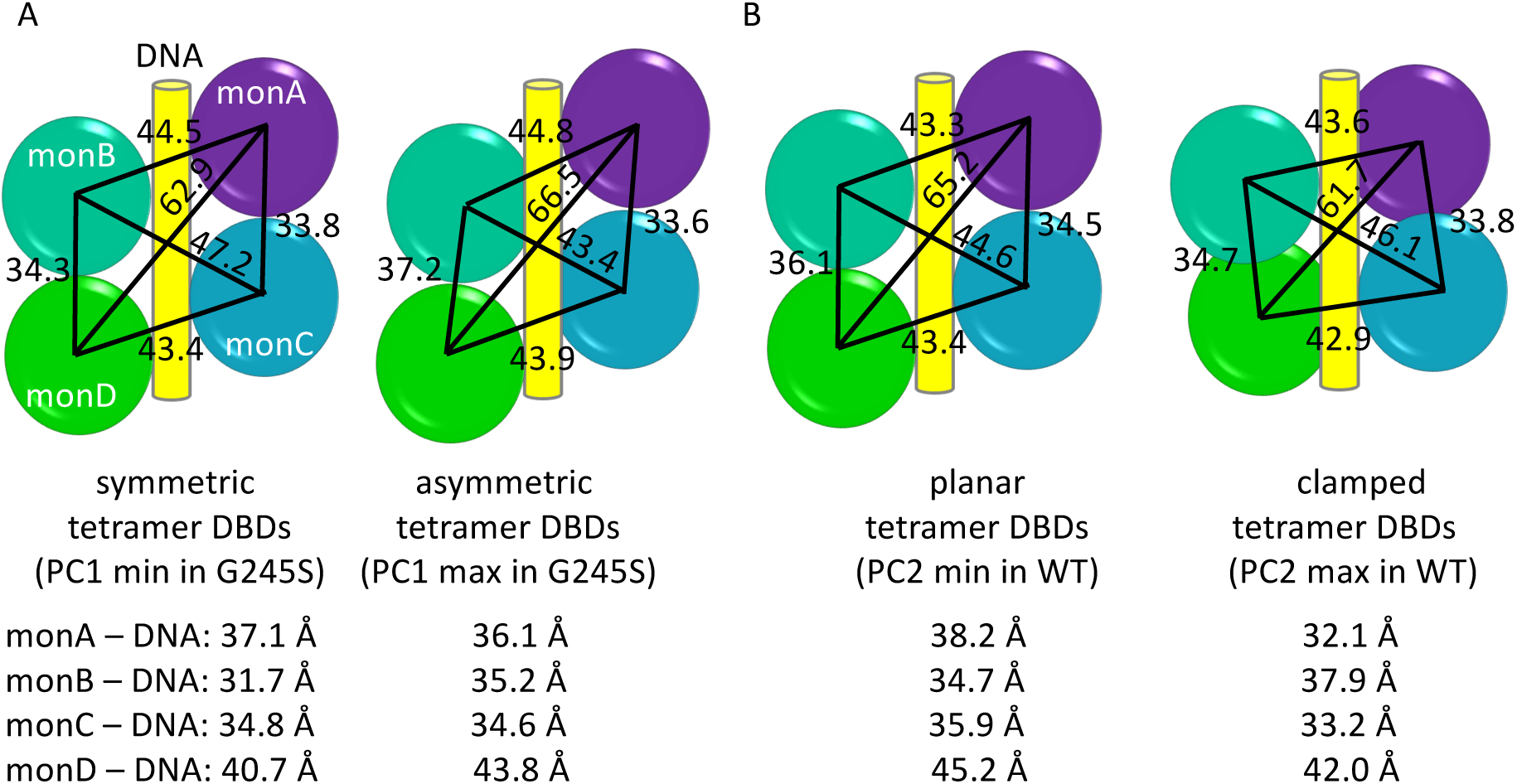
Quaternary binding modes of p53 DBD tetramer to DNA. A) Schemes for quaternary binding modes that correspond to minimum and maximum PC1 values for G245S system. B) Schemes for quaternary binding modes that correspond to minimum and maximum PC2 values for WT system. Distances between center of mass (COM) reported are all in Angstroms.

Additionally, the PCA plots in Figure 3 indicate that WT p53 samples conformations that correspond to high PC2 values not accessed by the two mutant systems. The transition from low to high PC2 values in the WT system translates into the DBDs of monomers A, C and D approaching DNA, the inner monomers B and C moving away from each other while the outer monomers A and D approach each other generating a tighter-bound, clamped tetramer conformation unique to WT p53 (Fig 4B). This clamped conformation has the H2 helices (residues 276-287) of monomers A and D slightly closer to the DNA groove and H2 helices of monomers B and C slightly further from the DNA groove resulting in a reduced solvent accessibility of the DNA response element (3888 Å^2^ compared to 3968 Å^2^ measured for WT PC2 minimum structure).

### DBD Dynamics

A previous study^18^ on the dynamics of the monomeric DBD of WT p53 and the Y220C mutant, utilizing extensive MD simulations and Markov state modeling (MSM), revealed that loops L1 and L6 are involved in the two slowest and most significant motions of the protein (Fig 1C). The computational models, validated by NMR relaxation experiments, established a set of features consisting of interacting pairs centered around loops L1 (residues 113–123) and L6 (residues 221– 230) to discretize the p53 DBD conformational space^18^. In this study, the previously established set of 24 features was used to compare the dynamics of the p53 DBD within DNA-bound full-length p53 tetramer systems to those of the monomeric p53 DBD system (Fig 1D). To directly compare the conformational landscapes of WT, Y220C mutant, and G245S mutant fl-p53 tetramer systems with the WT p53 DBD monomer system, the conformations explored by each system, represented by the 24 features, were used as input for time-lagged Independent Component Analysis (tICA)^30^. The resulting free energy landscapes are shown in Figure 5. Using tICA to jointly process the trajectories with the selected features ensures that the new coordinate space is consistent across all systems, making the conformational ensemble of each system directly comparable. tIC1 is described by L6 loop anchors and tIC2 is described by L1 loop anchor (Fig 1D). Since the tICs are ordered from slowest to fastest motions, the transitions involving L6 loop are slower than those involving L1 loop (Fig 1D). In the free energy landscape plots of Fig 5, smaller values of each tIC correspond to extended loop conformations, while larger tIC values correspond to recessed loop conformations.

**Figure 5.**
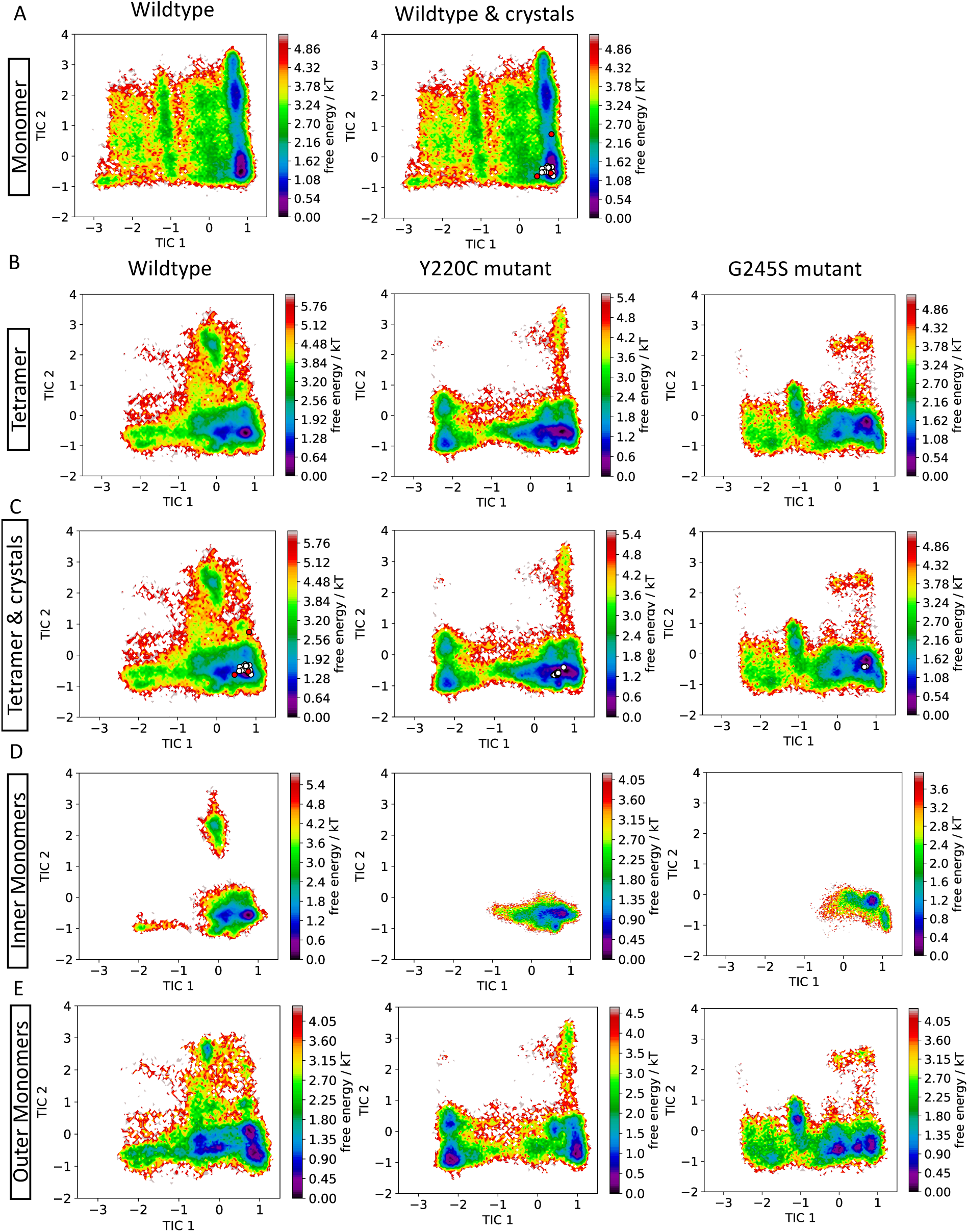
Free energy landscapes based on the 24 features jointly used as input for time-lagged Independent Components Analysis (tICA). tIC1 is described by L6 loop anchors, tIC2 is related to L1 loop anchor. For each tIC, smaller values of correspond to extended loop conformations, while larger values correspond to recessed loop conformations.

The WT p53 DBD monomer system has two preferred states that differ from each other in tIC2 coordinates corresponding to two different L1 loop conformations, recessed and extended (Fig 5A). In contrast, the preferred states for WT, Y220C mutant and G245S mutant fl-p53 tetramer systems occur at low tIC2 values, corresponding to an extended L1 loop conformation, but differ in their tIC1 values indicating different L6 loop conformations (Fig 5B). Overlay of the p53 crystal structures in the Protein Databank onto the free energy plots using the same coordinate space indicate an excellent overlap of most structures with the preferred state at tIC1= ∼1 and tIC2= -1 shared across all systems (Fig 5A, 5C). The only exceptions shown by the two fully overlapping red circles at tIC1=1 & tIC2=1 belong to the two outer monomers from DNA-bound tetrameric WT p53 crystal structure (pdbID 3TS8) which are known to have captured unique (recessed) L1 conformation (wildtype monomer & crystals plot in Fig 5A, wildtype tetramer plot in Fig 5C).

The fl-p53 tetramer forms a dimer of dimers and each p53 monomer binds to a pentamer repeat^31^. In the p53 DBD tetramer that consists of dimer1 (monomers A and B) and dimer2 (monomers C and D), two types of protein-protein interfaces are observed (Fig S13). The first is the monomer-monomer interface, located within each DBD dimer (A-B and C-D interfaces). This interface includes Zn-coordinating residues, the H1 helix, and parts of the L2 and L3 loops from both monomers. The second is the dimer-dimer interface (A-C and B-D interfaces), which involves the L1 loop (Thr123, Cys124) and L6 loop (Val225) from one monomer, along with the L3 loop (Ser166, Gln167) and the N-terminus of the DBD (residues 99-103) from the other monomer. The monomer-monomer interface directly interacts with DNA when DNA binds.

The CDKN1A (p21) response element, which was the first natural response element crystallized in complex with a p53 DBD tetramer, consists of 4 contiguous pentamer repeats. In the DNA-bound structure, the two inner p53 monomers (monomers B and C) bind to the two central pentamers of this DNA response element while the two outer p53 monomers (monomers A and D) bind to the two outer pentamers (Fig S13). The crystal structure of p53 tetramer bound to p21 response element (pdbID 3TS8) showed that the L1 loop adopts an extended conformation in the two inner monomers and a recessed conformation in the two outer monomers^10^. In all other p53 crystal structures examined (listed in Table S2, S3 & S4), L1 loop is found in an extended conformation. Because of this L1 loop variance, we explored whether the free energy landscape of the inner and outer monomers differ. Interestingly, in WT p53 as well as Y220C & G245S mutant systems, the free energy landscape sampled by the inner monomers (Fig 5D) is tightly centered around tIC1= ∼1 and tIC2= -1 and significantly more localized/constrained compared to that of the outer monomers (Fig 5E). Therefore, the location of the p53 monomer in the DNA-bound p53 tetramer proves to be important and governs the DBD dynamics it can sample. L1 loops of the two inner monomers are very close to the dimer-dimer interface in the tetramer and their more constrained conformational sampling is likely due to this proximity. The L1 loops of the two outer monomers, on the other hand, are far from dimer-dimer interfaces and sampled a larger conformational space.

To study the dynamics of the simulated p53 systems in more detail, we turned to Markov state models (MSMs). MSMs combine multiple MD simulations into a unified model of the protein’s conformational ensemble, capturing key thermodynamic and kinetic properties while preserving atomic-level details of the system. By applying MSMs to our extensive MD simulations, we sought to address sampling limitations and access a highly accurate description of the conformational ensemble, offering unprecedented insights into the systems under study.

### Markov state models centered on L6 loop

Markov state models constructed for all monomer data together using only the 17 features that include L6 anchor residues identify the presence of 4 metastable states for WT monomer, 6 metastable states for WT tetramer, 2 metastable states for Y220C tetramer and 4 metastable states for G245 tetramer systems (Fig 6) based on the relative separations of the slowest relaxation timescales in the implied timescale plots as detailed in Methods. The most populated metastable state across all systems, state R, has the highest tIC1 values and corresponds to the recessed L6 conformation (Table 1). This is also the only L6 conformation observed in all analyzed crystal structures (Fig 5A, 5C). Metastable state E with the lowest tIC1 values corresponds to the extended L6 conformation and has a stationary population of 18.0% in WT monomer, 8.0% in WT tetramer, 31.7% in Y220C tetramer and 16.2% in G245S tetramer (Fig 6, Table 1). The significantly reduced population of extended L6 conformation in WT tetramer compared to the WT monomer is likely because of the L6 loops of the two monomers that lie at the dimer interface in p53 tetramer structure. The stabilization of the extended L6 conformation in Y220C tetramer on the other hand is a direct effect of the mutation of residue 220, which resides at one end of L6 loop, on loop dynamics. Extended L6 conformation, which was identified for the first time in MD simulations of WT and Y220C mutant p53 DBD monomers in Barros *et al*^18^ and now observed in WT and mutant fl-p53 tetramers in this study, has the pocket typically targeted by small molecules closed but a novel pocket open on the opposite side of L6 loop (Fig S14). This novel cryptic pocket was only very recently captured bound to a ligand by the Shokat lab (pdbID 8DC6) and presents a unique opportunity for structure-based design of small molecule binders for Y220C mutant. It was not possible to include 8DC6 in Fig 5C as residues 115-122 are not resolved in the crystal structure. It is also notable that Y220C tetramer is unique in having no intermediate states between states R and E (Fig 6), in line with previous MSMs of Y220C DBD monomer^18^.

**Fig 6.**
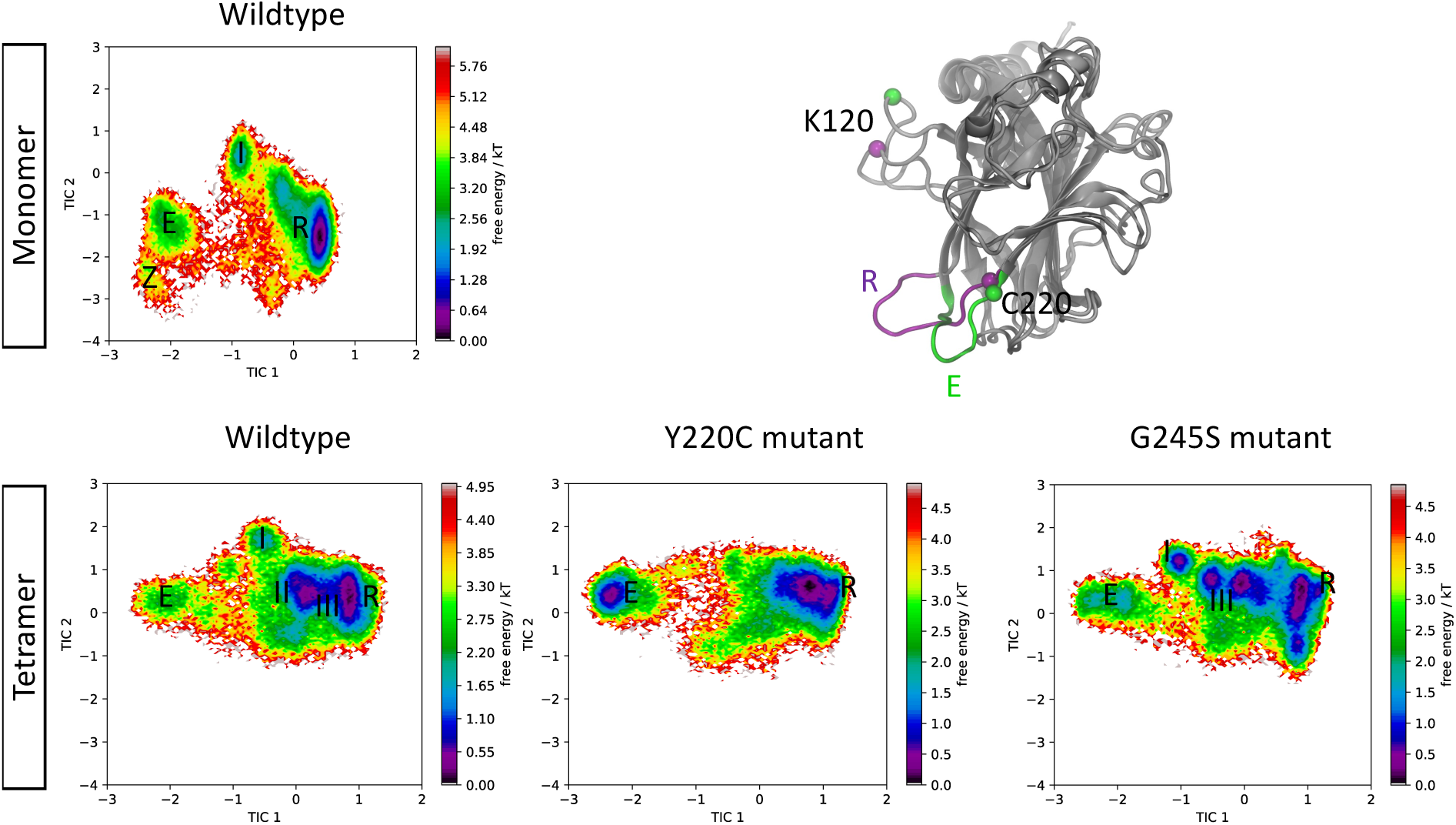
Markov state models centered on L6 loop. Free energy landscape of each system in terms of 17 L6 features are plotted with metastable states labeled. Low TIC1 values: Extended L6 conformation. High TIC1 values: Recessed L6 conformation. Extended (E) and recessed (R) L6 conformations in representative Y220C metastable state structures are depicted with purple and green ribbons, respectively. C220 and K120 C-alphas are represented with spheres.

**Table 1.** Equilibrium populations of metastable states in L6-centered MSMs.

| States | WT monomer | WT tetramer | Y220C tetramer | G245S tetramer |
| --- | --- | --- | --- | --- |
| E | 18.0% | 8.0% | 31.7% | 16.2% |
| R | 51.8% | 38.3% | 68.3% | 40.2% |
| I | 27.0% | 10.4% |  | 9.9% |
| II |  | 27.9% |  |  |
| III |  | 8.0% |  | 33.7% |
| Z | 3.1% |  |  |  |

### Markov state models centered on L1 loop

Wild-type p53 has intrinsic L1 loop flexibility indicated by available crystal and NMR structures^10^ and computational studies^14,32,33^. Our previous study on the p53 DBD^18^ showed how p53 mutations outside L1 loop can affect L1 dynamics. To monitor L1 dynamics in the fl-p53 tetrameric system, we constructed MSMs for all monomer data together and using only the previously established 7 features that include the L1 anchor residue Ser116. The MSMs (Fig 7) identify 5 metastable states for the WT DBD monomer, 4 metastable states for the WT tetramer and 3 metastable states each for the Y220C tetramer and G245 tetramer systems. Populations of these metastable states are listed in Table 2.

**Figure 7.**
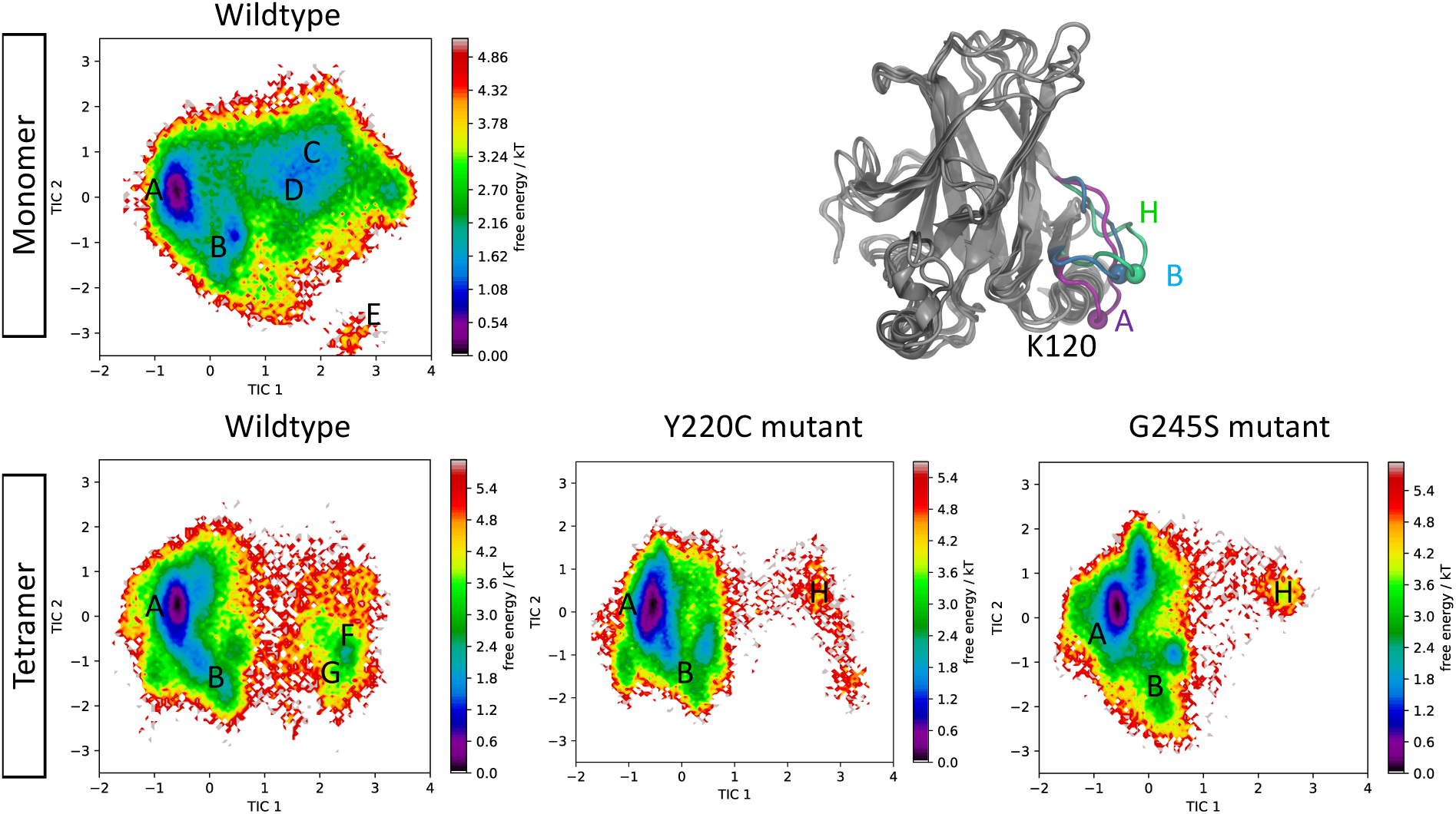
Markov state models centered on L1 loop. Free energy landscape of each system in terms of 7 L1 features are plotted and metastable states are labeled. Low TIC1 values: Extended L1 conformation, High TIC1 values: Recessed L1 conformation.

**Table 2.** Equilibrium populations of metastable states in L1-centered MSMs.

| States | WT monomer | WT tetramer | Y220C tetramer | G245S tetramer |
| --- | --- | --- | --- | --- |
| A | 41.8% | 65.3% | 62.6% | 58.6% |
| B | 21.1% | 18.9% | 14.0% | 16.0% |
| C | 13.1% |  |  |  |
| D | 9.6% |  |  |  |
| E | 14.4% |  |  |  |
| F |  | 10.5% |  |  |
| G |  | 5.2% |  |  |
| H |  |  | 23.4% | 25.4% |

State A is the most populated state across all systems and corresponds to the most extended conformation of L1 loop, which as discussed above is observed in almost all experimentally determined p53 structures. The population of state A is significantly larger in all tetrameric systems compared to the WT DBD monomer, likely due to the combined effect of the presence of DNA and tetrameric p53 organization. State B, with a 3-10 helix at the L1 loop, is also conserved across all systems with slight differences in their equilibrium populations. States C and D, identified in WT DBD monomer, are abrogated in all tetrameric systems. These two states have the L1 loop in recessed conformation and have a combined equilibrium population of 20%. The L6 loop is also in recessed conformation and interacts with the L1 loop via hydrogen bonds between Ser116 on L1 loop and Asp228 backbone N and Thr231 side chain in states C and D, respectively. State E, also only visited in the WT monomer, has an equilibrium population of 14.4% and shows the L1 loop adopting an excessively recessed conformation which lacks persistent L6 loop interactions. State F, with an equilibrium population of 10.5%, is solely observed in the WT tetramer and corresponds to a recessed L1 loop with a H-bond observed between Thr123 backbone N and Ser116 backbone carbonyl. State G in WT tetramer has an equilibrium population of 5.2% and has a recessed L1 loop in which residues 114-116 form a 3-10 helix. State H is observed only in the Y220C and G245S mutant tetramer systems, each having an equilibrium population of ∼25% and corresponding to recessed L1 loop conformations.

Using the FTMap^34^ algorithm to probe the surface of the representative structures of each state, the L1/S3 pocket is found ligandable/druggable in Y220C tetramer state A and G245S tetramer states A and H. In the L1/S3 pocket, residues Pro140, Cys141, Leu114, His115, Ser116, Cys124 interact with FTMap probes (Fig S15 A). We also observed an additional druggable region on the opposite side of the L1 loop, which can be called the L1/S3 back pocket, that binds to FTMap probes across all states and consists of residues Phe113, Leu114, His115, Trp146, Tyr128, Phe270, Asn131 (Fig S15 B).

### L1/S3 Pocket

In addition to exploring the dynamics of the L1 loop using long timescale MD simulations and MSMs, we also monitored how often the druggable pocket governed by the L1 loop was open. We previously discovered that the transiently open, cryptic L1/S3 pocket can be used for small molecule binding and eventual p53 reactivation^14–16^. There is now further experimental and structural evidence for druggability of the L1/S3 pocket^17,35,36^. This druggable pocket is present across all p53 mutants and offers a promising opportunity for structure-based discovery of p53 reactivators. Using the geometric criteria we established in Wassman et al^14^, the L1/S3 pocket is open 18.7%, 21.0% and 17.9% (averaging over all monomers of the tetramer) of the time in the MD simulations for WT, Y220C and G245S fl-p53 tetramer systems, respectively. In the inner monomers (B and C), the L1/S3 pocket is open 7 to 13% while in the outer monomers (A and D), the pocket is open 16 to 38% of the time (Table 3). It is worth noting that the difference in the L1/S3 pocket sampling frequencies between outer and inner monomers in this study is not as drastic as observed in our 2017 study^28^, which was likely biased by the initial starting structure (pdb:3TS8-based integrative model) due to limited sampling at 100 ns timescale. In summary, the L1/S3 pocket is accessible for small molecule binding in all monomers of the p53 tetramer with the outer monomers being more accessible than the inner monomers.

**Table 3.** Sampling frequency of open L1/S3 pocket observed throughout MD simulations using geometric criteria.

|  | **Monomer A** | **Monomer B** | **Monomer C** | **Monomer D** | **Mean** |
| --- | --- | --- | --- | --- | --- |
|  | **(%)** | **(%)** | **(%)** | **(%)** | **(%)** |
| WT | 16.3 | 8.5 | 12.9 | 37.0 | 18.7 |
| Y220C | 29.0 | 10.3 | 7.2 | 37.6 | 21.0 |
| G245S | 32.1 | 9.5 | 12.1 | 18.0 | 17.9 |

We also monitored the solvent accessibility of the C124, C135 and C141 cysteine triad sidechains at the L1/S3 pocket, which were found coordinated to arsenic in the crystal structures that elucidated how FDA-approved arsenic trioxide (ATO) binds and reactivates p53^17^. Table 3 indicates that the average solvent accessible surface area (SASA) of the C124 sidechain is in the range of 2-8 Å^2^ for WT p53 tetramer as well as the two cancer mutants. The average C124 sidechain SASA is noticeably higher for monomer C than other monomers in all systems. For comparison, C124 sidechain SASA in the p53 DBD crystal structure 1TSR chain B is 0.43 Å^2^, indicative of a fully buried C124. By calculating the frequency of MD frames with a C124 sidechain SASA above 5 Å^2^ (a selected cutoff to define C124 solvent exposure), we found C124 to be solvent-exposed 10-70% of the time in the simulations (Table 4). Again, monomer C has significantly higher C124 exposure in all systems.

**Table 4.**
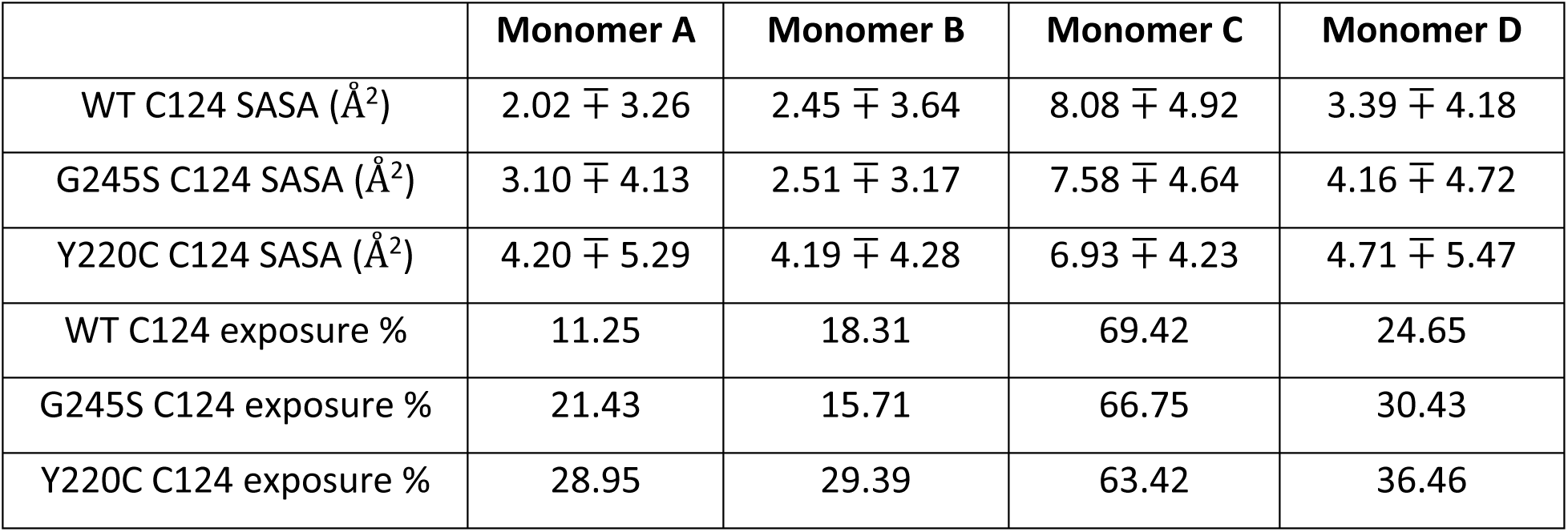
Solvent-accessibility of C124 sidechain at L1/S3 pocket (average ∓ standard deviation) and percentage of MD frames with C124 SASA > 5 Å^2^.

The average C141 sidechain SASA ranges between 0.5 and 1.12 Å^2^ for all monomers across all systems (Table S5). The average C135 sidechain SASA is even lower, ranging between 0.1 and 0.5 Å^2^ (Table S6). Using the same SASA criteria for solvent exposure, C141 and C135 are solvent exposed only in 0.8-3.7% and 0-1.3% of the frames in the MD simulations, respectively. Thus, among the cysteine triad at the L1/S3 pocket, C124 is the most solvent exposed cysteine and C135 the least exposed. Additionally, in most cases, C124 is more exposed in the cancer mutants than the WT. For comparison, we also calculated the solvent accessibility of the remaining 4 cysteines that are not zinc-coordinating in DBD of WT p53 (Table S7). Based on MD-averaged solvent exposure frequencies in the DNA-bound WT fl-p53 tetramer, C277 ∼ C182 > C229 > C124 > C275 ∼ C141 > C135. This order can provide insight into the availability of p53 cysteines for covalent attachment of small molecules when DNA-bound fl-p53 tetramer is in solution.

## DISCUSSION

All-atom microsecond-timescale MD simulations in this study aim to elucidate the dynamics of the functional form of p53 under physiological conditions (*in situ*). Assembling the crystal structures of isolated p53 domains, we constructed an integrated model of the DNA-bound full-length p53 (fl-p53) tetramer in explicit solvent, for the WT p53 as well as the G245S and Y220C mutants. Using the Anton2 supercomputer, we explored their long-timescale dynamics. This analysis allowed us to observe, in atomic detail, how different domains of fl-p53 interact with one another, with other p53 monomers in the tetramer, and with DNA. Additionally, it enabled a direct comparison of the G245S and Y220C cancer mutants with WT p53 in the context of a DNA-bound full-length tetramer. Our findings align with a growing body of work showing that simulating proteins within their full physiological context yields more accurate and often unanticipated insights−such as hinge-mediated allostery in a nuclear receptor heterodimer^37^, cryptic epitope exposure during breathing and tilting motions of influenza glycoproteins^38^, and cholesterol-sensitive structural rearrangements in a sodium channel^39^−highlighting the unique advantage of *in situ* simulations for capturing cryptic, functionally important dynamics that are inaccessible to isolated proteins.

Our initial fl-p53 tetramer models had fully extended CTD and NTD domains. However, after conducting nanosecond-timescale MD simulations^28^, we previously observed interactions between the CTD and DNA. In this study, using microsecond-timescale MD, we identified new, dynamic interactions between the NTD and p53 DBD. The DNA bending observed aligns quantitatively with experimentally reported DNA bending angles. The consistency between multiple simulation observables and experimental data strengthens confidence in the novel insights from these MD simulations, particularly those that cannot be easily captured through experimental methods.

Our previous nanosecond-timescale MD studies have shown in atomic detail that neither a p53 cancer mutant in complex with response element DNA nor WT p53 in complex with nonspecific DNA sequence can retain optimal quaternary binding mode of the p53 tetramer to DNA^28,29^. A stable p53 and a specific DNA are both required for optimal quaternary organization. In this study, when we monitor the dynamics of the p53 DBD tetramer organization around DNA with PCA analysis (Fig 3), WT p53 DBD tetramer samples a tightly clamped organization that is not accessed by the two cancer mutants and is different from the starting crystal structure (Fig 3, 4B). This tightly clamped WT p53 DBD tetramer conformation might be the real specific DNA-binding mode of p53 (or an intermediate state towards that state) that is not exhibited in X-ray crystal structures of p53-DNA complex but can only be accessed “in the presence of all p53 domains and after dynamics” as suggested by Ouaray et al 2016^33^.

PCA also captured an asymmetric organization of the DBD tetramer sampled only by the G245S mutant (Fig 3, 4A, S12). The CTD of G245S monomer B has almost no contacts with DNA (Fig S7) and PCA results in Fig 4A showed that the DBDs of monomers B and D of this mutant moved away from DNA as captured by the conformational variation along PC1. We hypothesize that these are likely signs of initial dissociation of G245S tetramer from DNA and indicate its non-optimal binding. We also observed an interaction between Ser183 in monomer C and Asp186 in monomer B in the highest PC1 conformation. Notably, phosphorylation of Ser183 has been reported to affect the DNA binding cooperativity of p53^40^. Additionally, phosphorylation of p53 Ser183 by

Aurora B kinase marks it for proteasomal degradation^41^. Given that p53 cancer mutants are known to accumulate in cells, it is plausible that certain mutants, such as G245S, may adopt a different tetrameric organization that conceals phosphorylation sites like Ser183, potentially delaying or preventing the proteasomal degradation of these cancer mutants.

In addition to studying the behavior of global features in the fl-p53 tetramer systems, the dynamics of the cryptic L1/S3 druggable pocket and the cysteine triad are of particular interest. MSM analysis of the L1 loop reveals that equilibrium populations of the extended L1 loop are larger, while those of the recessed L1 loop are smaller in DNA-bound tetramer systems compared to the p53 DBD monomer system. Furthermore, the metastable states involving L1-L6 loop interactions, observed in the p53 DBD monomer, are absent in the tetramer systems.

These population changes in the MSMs are likely asserted by the dimer-dimer interfaces of the tetramer complex. Using geometric criteria, we found that the L1/S3 pocket is open about 20% of the time across all fl-p53 tetramer system simulations, with the outer monomers having the L1/S3 pocket open more often compared to the inner monomers. Among the cysteine triad at the L1/S3 pocket, our analysis indicated C124 as the most solvent-exposed and C135 as the least solvent-exposed. C124 solvent exposure is higher in cancer mutants than WT but doesn’t differ in inner and outer monomers, in contrast to L1/S3 pocket opening frequency.

It is worth noting that zinc is coordinated to the protein and is not allowed to dissociate in our MD simulations. As a result, our simulations do not capture the structural effects of zinc loss observed to varying degrees for different p53 cancer mutants. Despite this limitation, the observables calculated from these microsecond-timescale MD simulations align well with experimentally reported values. We hope this study offers valuable insights into the physiologically relevant DNA-bound fl-p53 tetramer, which has eluded complete structural characterization, and provides some answers at full-atom detail. The structural ensembles generated in this study may also open new avenues for further exploration of this fascinating and important tumor suppressor protein.

## Acknowledgement

We thank Lorenzo Casalino for helpful discussions and feedback on figure generation. Anton 2 computer time was provided by the Pittsburgh Supercomputing Center (PSC) through Grant R01GM116961 from the National Institutes of Health; and an allocation under PSCA18040P to REA. The Anton 2 machine at PSC was made available by D.E. Shaw Research. This work was also supported by NIH R01 GM 132826.

## Data Availability

All fl-p53 tetramer MD trajectories & topology files as well as pdb files of last MD frames are available in the Amaro lab website at https://amarolab.ucsd.edu/files/p53/odemir/fl_p53_Anton. Similarly, pdb files of the structures that correspond to lowest & highest PC1 & PC2 values in PCA analysis and representative structures for Markov state models as well as their FTMap results are available for download.

## METHODS

### MD System Setup

Our DNA-bound fl-p53 tetramer model of WT p53 after ∼100 ns MD simulation from Demir et al 2017 study^28^ was used as the starting conformation, after trimming 8-9 nucleotides from each side of the DNA to reduce system size. For each cancer mutant, the relevant residues were mutated manually from the WT system. AMBER FF14SB force field^42^ parameters for protein, BSC1 force field^43^ for DNA, ZAFF^44^ parameters for zinc ion and its coordinating residues, and TIP3P^45^ parameters for waters were used. Na ions were added to neutralize the system.

Solvated WT system has 491,237 atoms (27,313 atoms for the protein and DNA only). Y220C system has 490,843 atoms (27,273 atoms for the protein and DNA only). G245S system has 491,253 atoms (27,329 atoms for the protein and DNA only).

### MD System Preparation

MD simulations with Amber18 suite^46^ were used to relax the system before starting the Anton MD production runs. First, a multi-step restrained minimization protocol followed by an unrestrained minimization was used to minimize each system. Then, each system was heated from 0 K to 310 K keeping protein heavy atoms restrained and relaxed with multi-step 250 ps MD simulations slowly decreasing the protein heavy atom restraints. A final unrestrained 10 ns MD simulation at 310 K prepared each system for Anton2 runs. Additional replicates of each system were generated by simulating for 2 additional nanoseconds to randomize the input velocities. MD simulations on Anton2 were run as an NPT ensemble using Anton’s multigrator^47^ at 300 K, 1 bar, and with a 2.5-fs time step and particle-mesh Ewald (PME) electrostatic approximations. For each system, 3 independent MD replicas of 7.5 µs each were performed without any constraints and system coordinates were saved with 0.24 ns intervals (corresponding to 93,750 time points for each of WT, G245S and Y220C systems).

### RMSD and RMSF Analysis

Root-mean-square-deviation (RMSD) and root-mean-square-fluctuation (RMSF) values were calculated using the cpptraj^48^ module of Amber18 suite. MD trajectories were first aligned to the starting MD structure with respect to the C-alpha atoms of the p53 tetramer DBDs.

### Secondary Structure Analysis and Principal Component Analysis (PCA)

Secondary structure analysis and principal component analysis were performed with mdtraj software^49^. PC1 and PC2 explain 17% and 13% of the variance, respectively.

### Markov State Model Construction

PyEMMA software^50^ was used to process simulation data and build Markov state models. Using the 24 features identified for p53 DBD in Barros et al^18^, we generated free energy landscapes by jointly processing all p53 variant trajectories as input for time-lagged Independent Components Analysis (tICA) with a tICA lag time of 6 ns^30^. L6-centered Markov state models were built using the 17 features that are centered at L6 loop. L1-centered Markov state models were built using the 7 features that are centered at L1 loop. Discretization was achieved by k-means clustering with k=150 for each system (WT monomer, WT tetramer, Y220C, G245S) separately, and accuracy of the models were confirmed by implied time scale plots and Chapman-Kolmogorov tests (Fig S16-S23). L6-centered and L1-centered Markov state models were both constructed with MSM lag time of 6 ns each.

### Cys Sidechain SASA

Cysteine sidechain SASA values were calculated using tcl scripts in VMD^51^ version 1.9.4a55 using a stride of 5.

