## Supporting Information for "Towards *In Situ* Dynamics of DNA-bound Full-Length p53 Tetramer"

**SUPPLEMENTAL INFORMATION**

**Özlem Demir[1,2], Emilia P. Barros[1,3], Rommie E. Amaro[4]**

**1** Department of Chemistry and Biochemistry, University of California, San Diego, La Jolla, California 92093, USA

**2** Current address: Novartis Biomedical Research, San Diego, California 92121 USA

**3** Current address: MSD R&D Innovation Centre, 120 Moorgate, London, U.K.

**4** Department of Molecular Biology, University of California, San Diego, La Jolla, California 92093, USA

**Supplementary Figure & Table Captions:**

Figure S1. RMSD analysis of the backbone atoms of all four DBDs in p53 tetramer systems.

Figure S2. Secondary structure analysis of DBD throughout MD simulations

Figure S3. RMSF analysis of C-alpha atoms in p53 tetramer DBDs.

Figure S4. A zoomed view on DBD residues of the RMSF analysis of C-alpha atoms in p53 tetramer DBDs.

Figure S5. DNA angle (in degrees) in fl-tetramer MD simulations as measured by the angle between specified atoms at the two termini of DNA and a specified atom at the center of DNA. All 3 MD replicas for each system are concatenated for plotting. DNA bending is calculated by subtracting the mean value obtained from the plots from 180 degrees.

Figure S6. Radius of gyration of fl-p53 tetramer in MD simulations (excluding DNA) calculated by VMD. X-axis is frame numbers while y-axis is radius of gyration in Å.

Figure S7. CTD interactions (per residue) with DNA in fl-p53 tetramer MD simulations. For each MD frame, the distance is calculated between each pair's closest heavy atoms (pairwise distances between p53 CTD residues and DNA nucleotides). It is counted as a contact if pairwise distance is less than 3.5 Å and the data is collected throughout the MD trajectories for the plot.

Table S1. Number of contacting residues between specified NTD and all DBD domains. (VMD analysis results) The distance is calculated between each pair's closest heavy atoms (pairwise distances between p53 DBD residues and NTD residues), and it's counted as a contact if that distance is less than 4.5 Å. Only the interactions that are observed more than 10% of the MD trajectory are listed.

Figure S8. WT DBD interactions (per residue) with NTD. The distance is calculated between each pair's closest heavy atoms (pairwise distances between p53 DBD residues and NTD residues), and it's counted as a contact if that distance is less than 3.5 Å.

Figure S9. Y220C DBD interactions (mean number of contacts per residue) with NTD in simulations. The distance is calculated between each pair's closest heavy atoms (pairwise distances between p53 DBD residues and NTD residues), and it's counted as a contact if that distance is less than 3.5 Å.

Figure S10. G245S DBD interactions (mean number of contacts per residue) with NTD in simulations. The distance is calculated between each pair's closest heavy atoms (pairwise distances between p53 DBD residues and NTD residues), and it's counted as a contact if that distance is less than 3.5 Å.

Figure S11. NTD interactions (mean number of contacts per residue) with DBD in simulations. The distance is calculated between each pair's closest heavy atoms (pairwise distances between p53 DBD residues and NTD residues), and it's counted as a contact if that distance is less than 3.5 Å.

Figure S12. Quaternary binding modes of p53 G245S mutant DBD tetramer to DNA observed in MD simulations. A) G245S conformations with lowest and highest PC1 values. B) Time evolution of the distance between D186 CG atom of monomer B and S183 OG atom of monomer C in MD simulations of G245S mutant.

Figure S13. The p21 (CDKN1A) response element consists of 4 contiguous pentamer repeats and the two inner p53 monomers bind the two central pentamers of this response element while the two outer p53 monomers bind the two outer pentamers

Table S2. The list of WT p53 crystal structures examined & used for comparison in conformational landscapes of Figure 5.

Table S3. The list of Y220C p53 crystal structures examined & used for comparison in conformational landscapes of Figure 5.

Table S4. The list of G245S p53 crystal structures examined & used for comparison in conformational landscapes of Figure 5.

Figure S14. Druggable pockets identified in representative structures of L6 loop MSM metastable states of Y220C. FTMap probes are depicted as cyan spheres. In recessed (R) conformation, L6 loop is separate from the loop of T150 and a ligandable pocket exists between the two loops. In extended (E) conformation, L6 loop approaches the loop of T150 closing the first pocket and opens a novel cryptic pocket on the opposite side of the L6 loop.

Figure S15. Druggable pockets identified in representative structures of L1 loop MSM metastable states. A) G245S metastable state A with open L1/S3 pocket conformation depicted as purple surface representation.

B) G245S metastable state H with open L1/S3 back pocket conformation depicted as green surface representation. FTMap probes are depicted as cyan spheres.

Table S5. Solvent-accessibility of C141 sidechain at L1/S3 pocket (average $\mp$standard deviation) and percentage of MD frames with C141 sidechain SASA > 5 $Å^2$

Table S6. Solvent-accessibility of C135 sidechain at L1/S3 pocket (average $\mp$ standard deviation) and percentage of MD frames with C135 sidechain SASA > 5 $Å^2$

Table S7. Solvent-accessibility of C182, C229, C275 and C277 sidechain (average ∓ standard deviation) in WT fl-p53 tetramer and percentage of MD frames with cysteine sidechain SASA > 5 $Å^2$

Figure S16. L6-centered MSM model validation analysis for WT monomer DBD. A. Implied timescale plot, B. Chapman-Kolmogorov test.

Figure S17. L6-centered MSM model validation analysis for WT tetramer. A. Implied timescale plot, B. Chapman-Kolmogorov test.

Figure S18. L6-centered MSM model validation analysis for Y220C tetramer. A. Implied timescale plot, B. Chapman-Kolmogorov test.

Figure S19. L6-centered MSM model validation analysis for G245S tetramer. A. Implied timescale plot, B. Chapman-Kolmogorov test.

Figure S20. L1-centered MSM model validation analysis for WT monomer DBD. A. Implied timescale plot, B. Chapman-Kolmogorov test.

Figure S21. L1-centered MSM model validation analysis for WT tetramer. A. Implied timescale plot, B. Chapman-Kolmogorov test.

Figure S22. L1-centered MSM model validation analysis for Y220C tetramer. A. Implied timescale plot, B. Chapman-Kolmogorov test.

Figure S23. L1-centered MSM model validation analysis for G245S tetramer. A. Implied timescale plot, B. Chapman-Kolmogorov test.

Figure S1

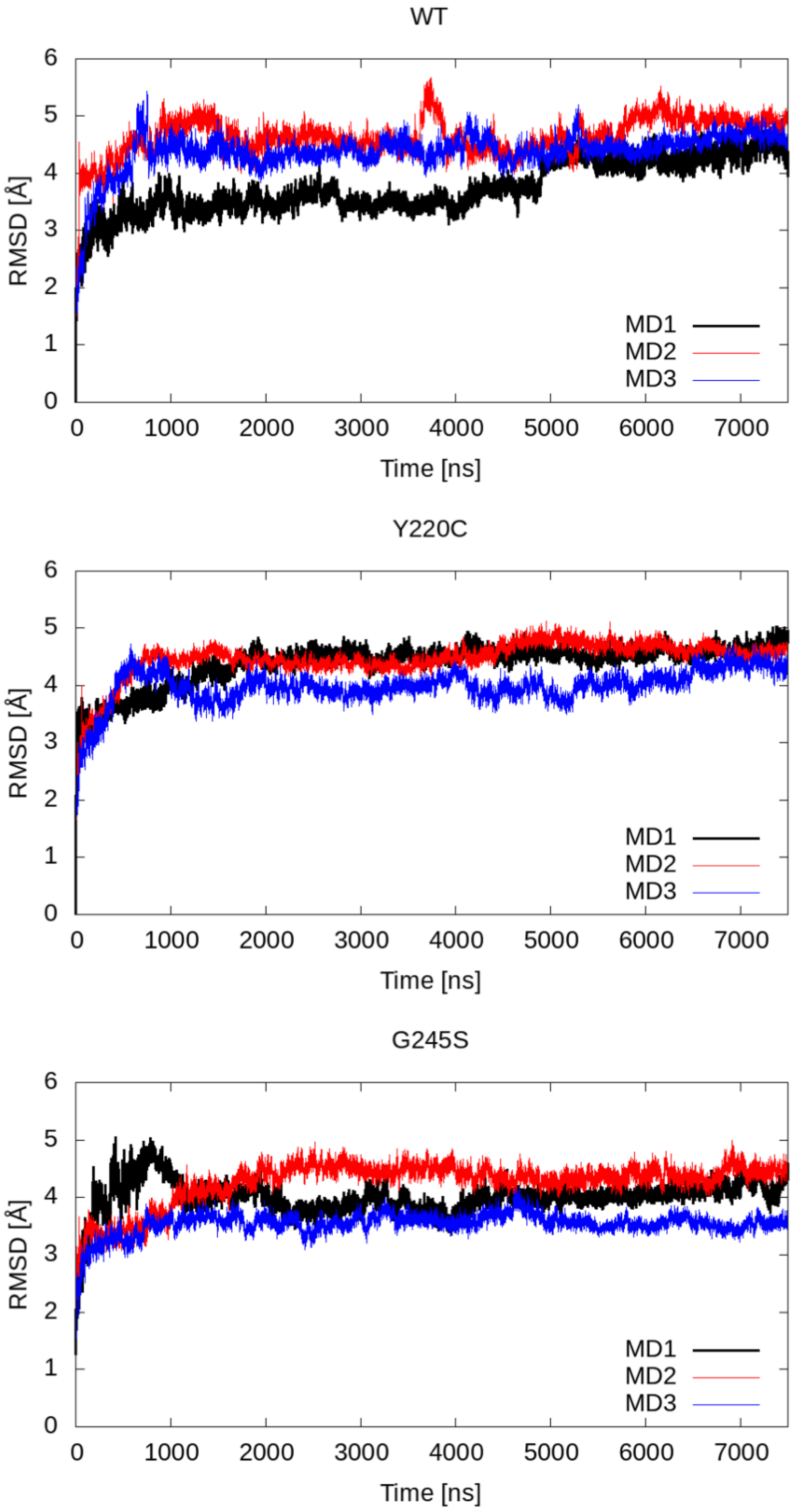

Figure S2

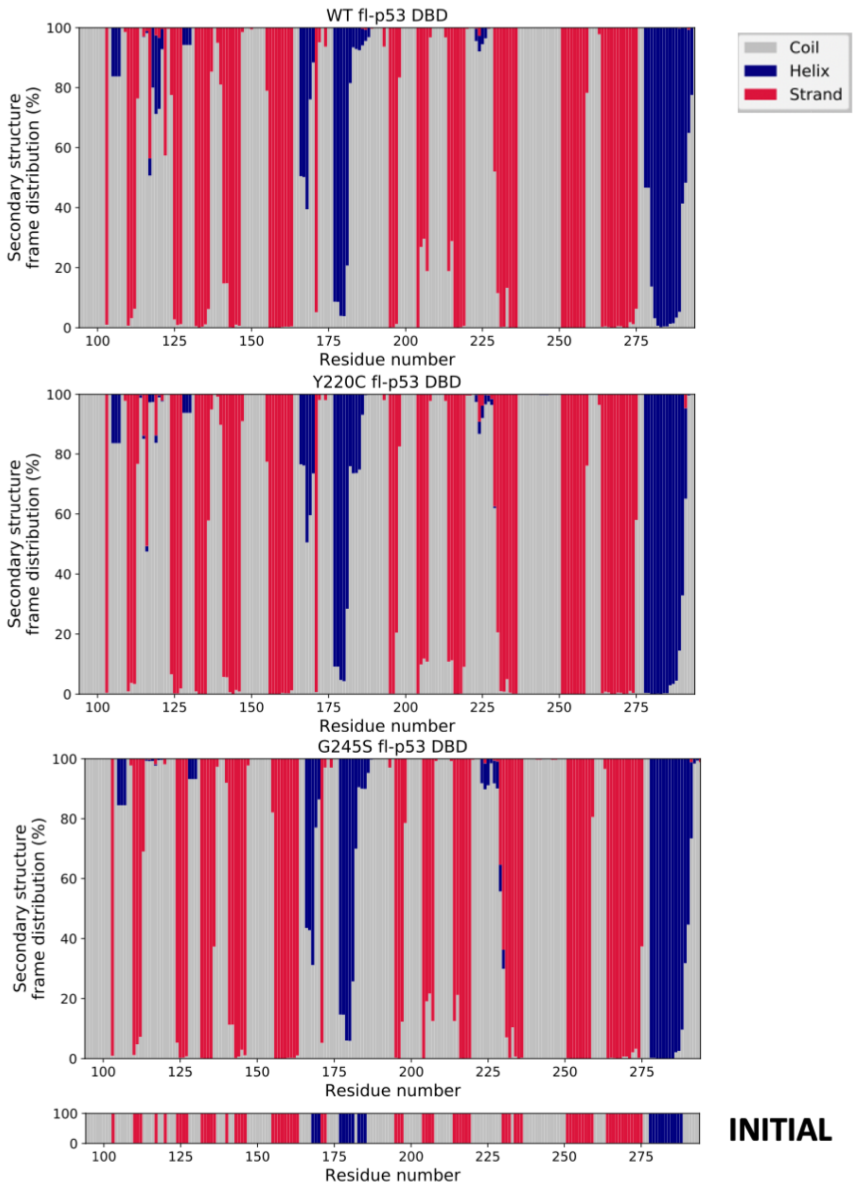
WT fl-p53 DBD
Y220C fl-p53 DBD
G245S fl-p53 DBD
Secondary structure frame distribution (%)
Residue number
100
125
150
175
200
225
250
275
0
20
40
60
80
100
Coil
Helix
Strand
INITIAL

Figure S3

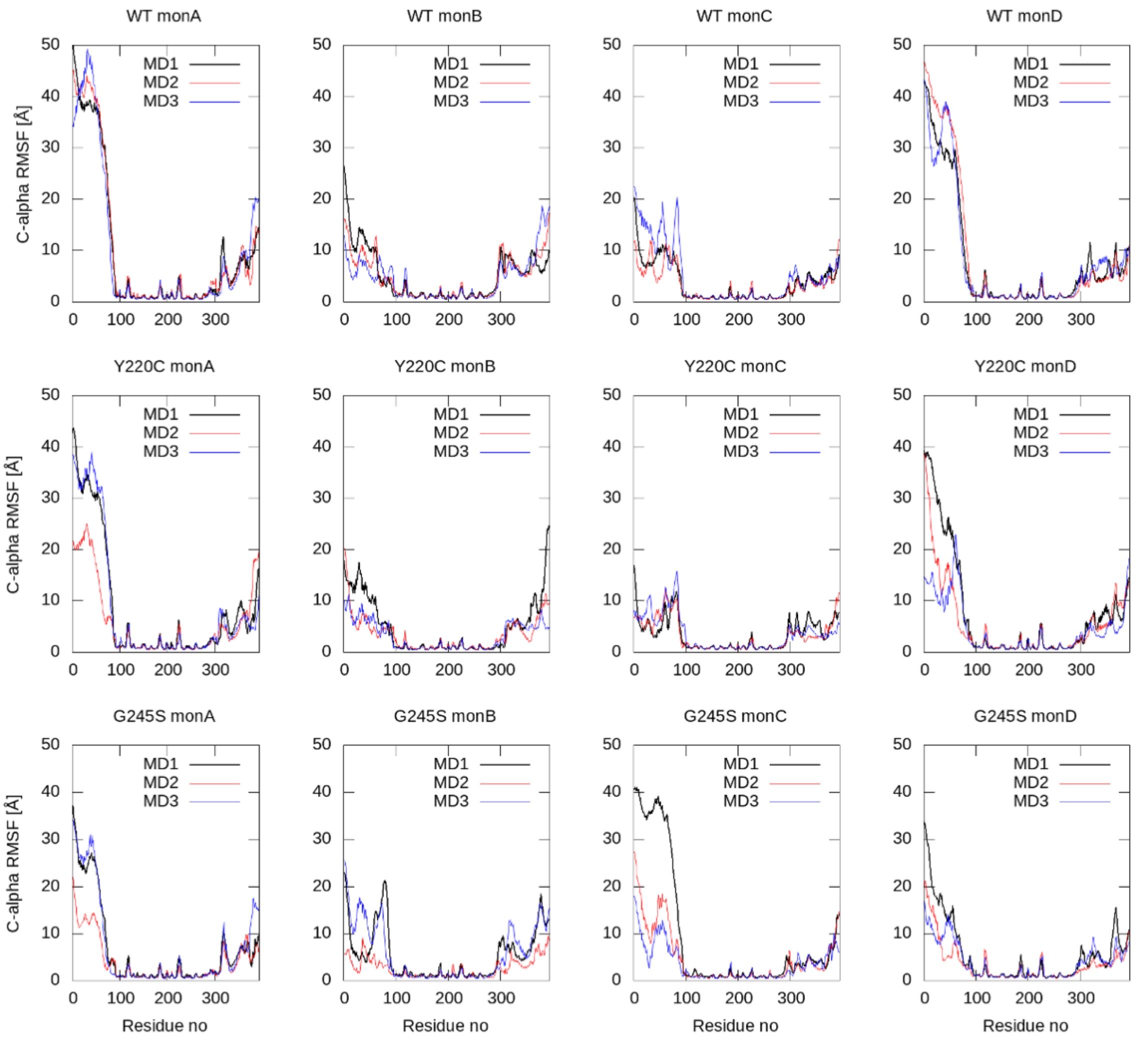
WT monA
WT monB
WT monC
WT monD
Y220C monA
Y220C monB
Y220C monC
Y220C monD
G245S monA
G245S monB
G245S monC
G245S monD
MD1
MD2
MD3
C-alpha RMSF [Å]
Residue no
0
100
200
300
10
20
30
40
50

Figure S4

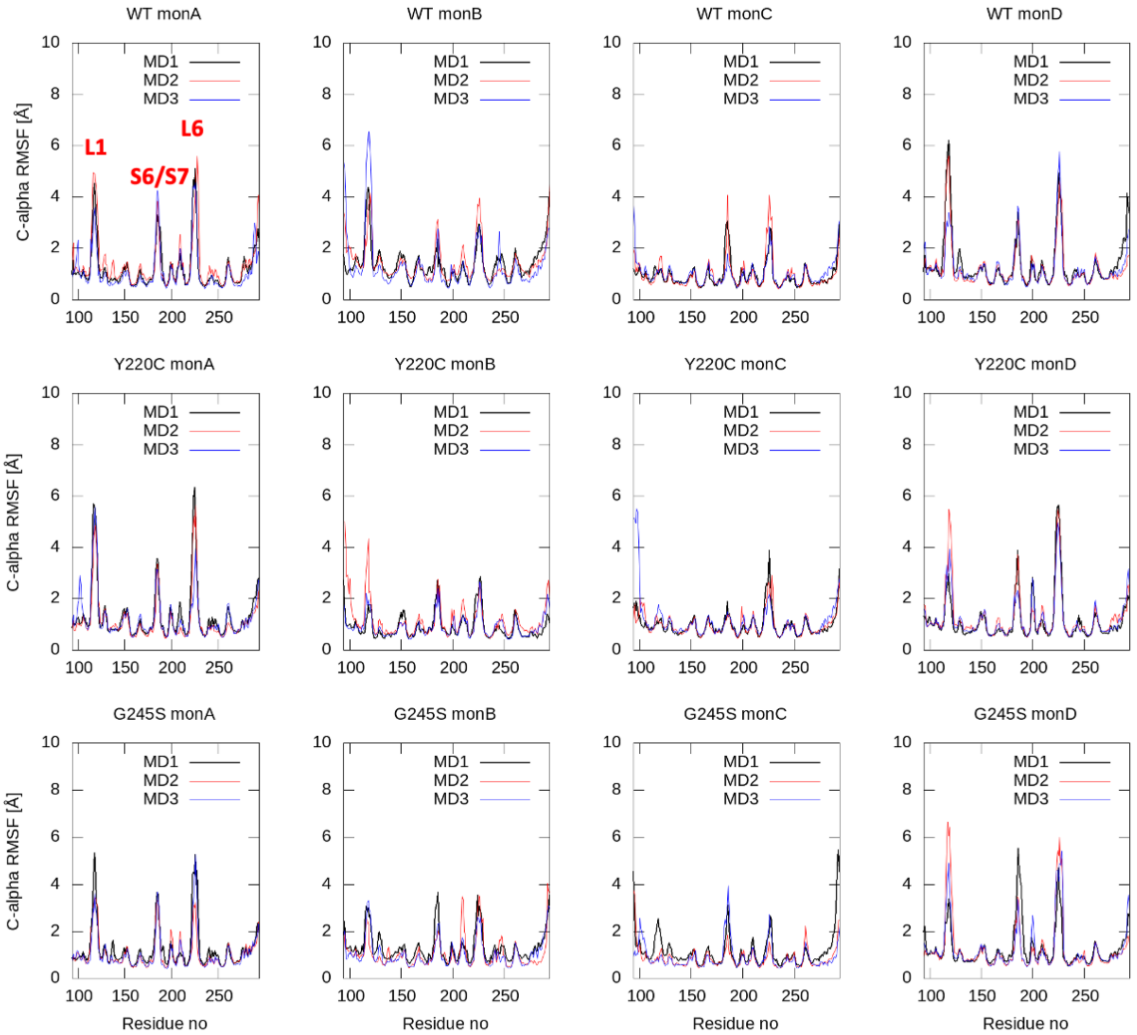
WT monA
WT monB
WT monC
WT monD
Y220C monA
Y220C monB
Y220C monC
Y220C monD
G245S monA
G245S monB
G245S monC
G245S monD
MD1
MD2
MD3
L1
S6/S7
L6
C-alpha RMSF [Å]
Residue no

Figure S5

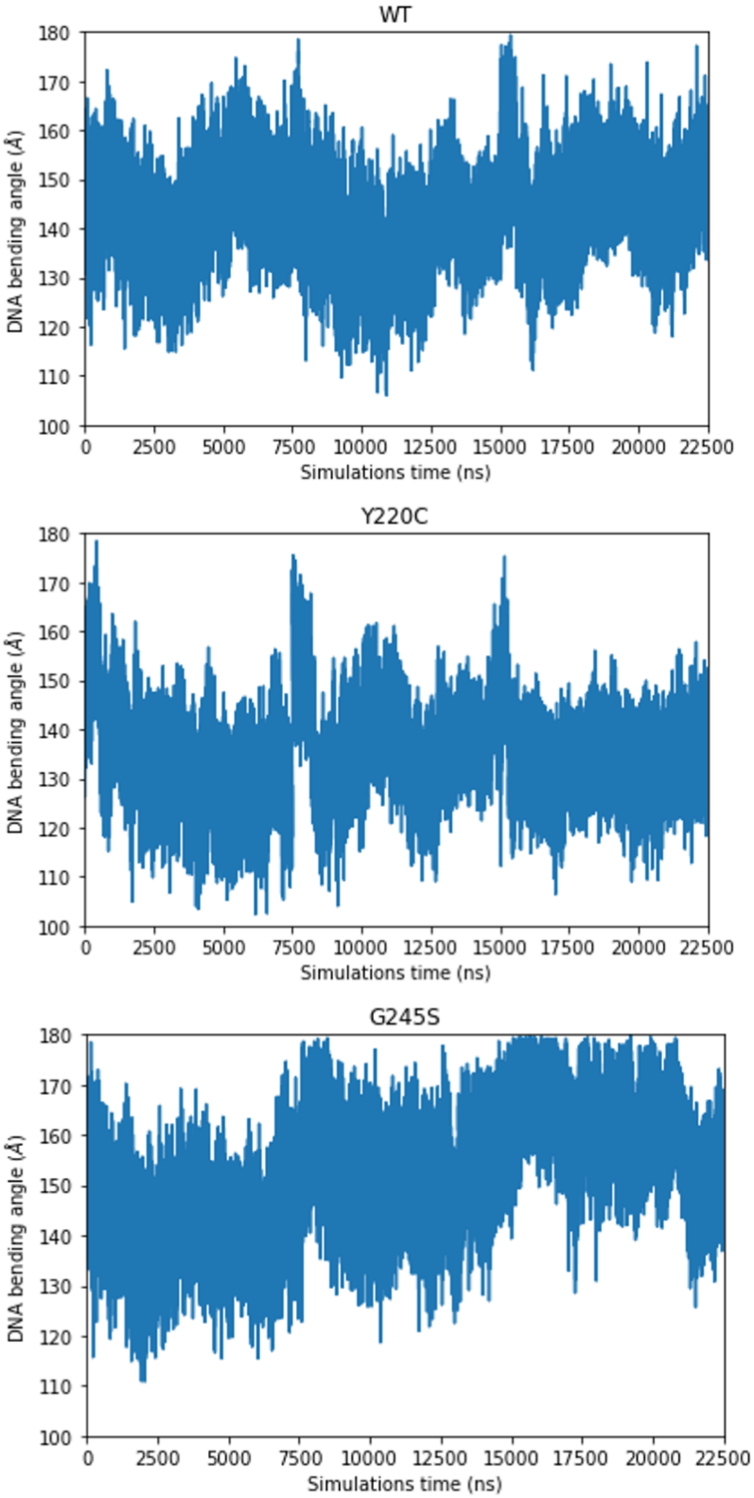
WT
DNA bending angle (Å)
Simulations time (ns)
Y220C
DNA bending angle (Å)
Simulations time (ns)
G245S
DNA bending angle (Å)
Simulations time (ns)

Figure S6

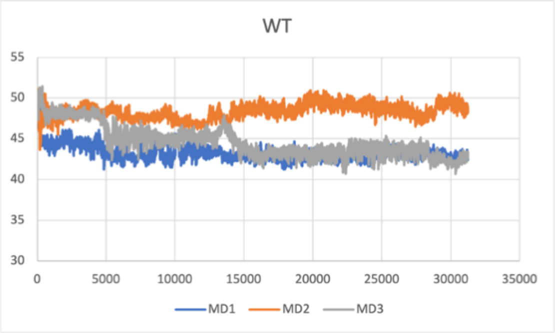
WT
55
50
45
40
35
30
0
5000
10000
15000
20000
25000
30000
35000
MD1
MD2
MD3

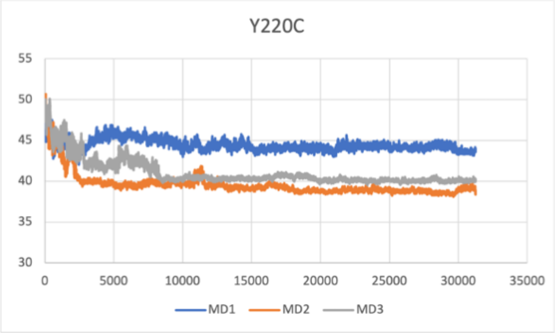
Y220C
55
50
45
40
35
30
0
5000
10000
15000
20000
25000
30000
35000
MD1
MD2
MD3

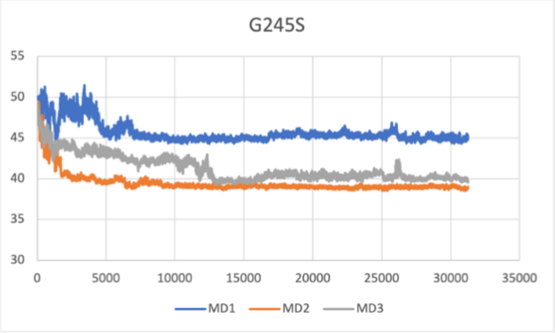
G245S
55
50
45
40
35
30
0
5000
10000
15000
20000
25000
30000
35000
MD1
MD2
MD3

Figure S7

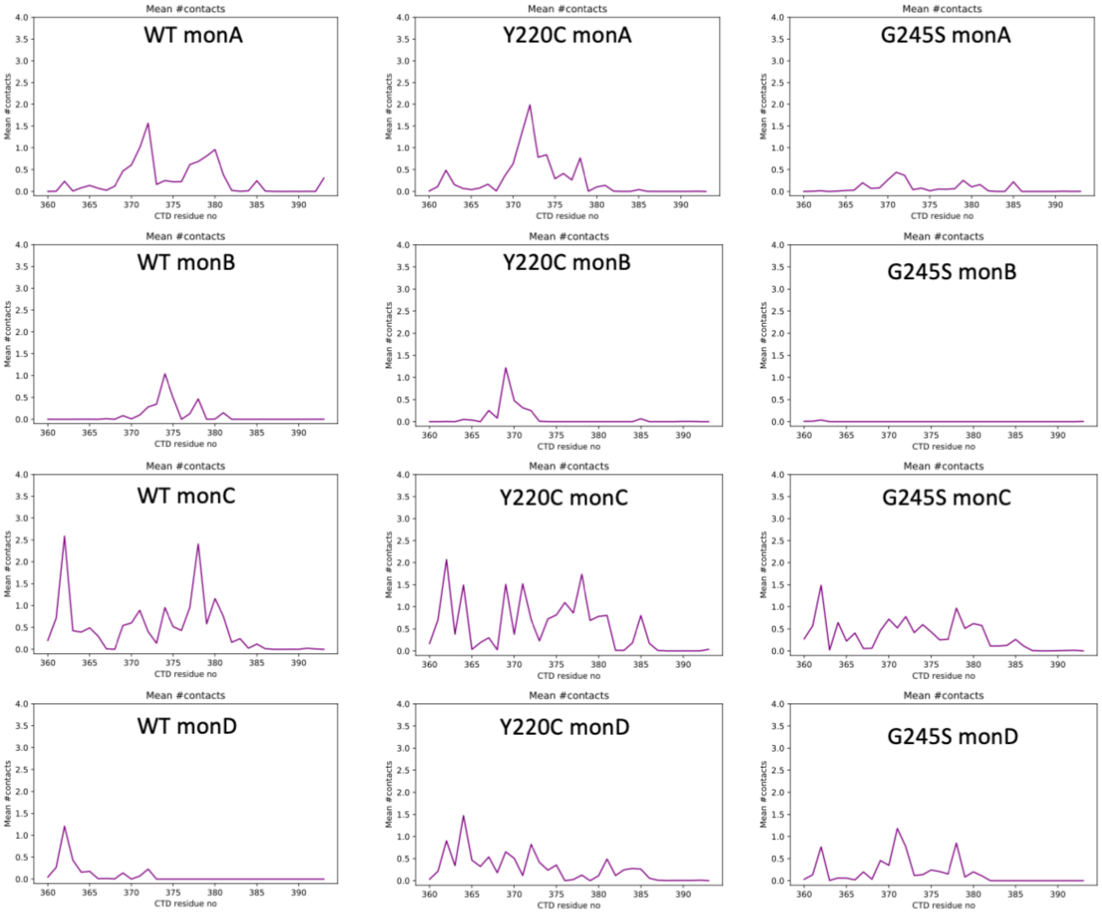


Sequence of p53 CTD:

360 – GSRAHSSHLK SKKGQSTSRH KKLMFKTEGP DSD -393

Table S1. Number of contacting residues between specified NTD and all DBD domains. (VMD analysis results) The distance is calculated between each pair's closest heavy atoms (pairwise distances between p53 DBD residues and NTD residues), and it's counted as a contact if that distance is less than 4.5 Å. Only the interactions that are observed more than 10% of the MD trajectory are listed.

| | Number of contacting residues (NTD + DBD) | Number of contacting residues in NTD | Number of contacting residues in DBDs |
|---|---|---|---|
| WT NTD-monA | 36 | 9 | 27 |
| WT NTD-monB | 184 | 78 | 106 |
| WT NTD-monC | 143 | 73 | 70 |
| WT NTD-monD | 58 | 19 | 39 |
| WT allNTDs | 400 | | |
| Y220C NTD-monA | 167 | 64 | 103 |
| Y220C NTD-monB | 186 | 82 | 104 |
| Y220C NTD-monC | 131 | 72 | 95 |
| Y220C NTD-monD | 132 | 57 | 75 |
| Y220C allNTDs | 563 | | |
| G245S NTD-monA | 138 | 59 | 79 |
| G245S NTD-monB | 189 | 83 | 106 |
| G245S NTD-monC | 120 | 50 | 70 |
| G245S NTD-monD | 179 | 80 | 99 |
| G245S allNTDs | 551 | | |

Figure S8

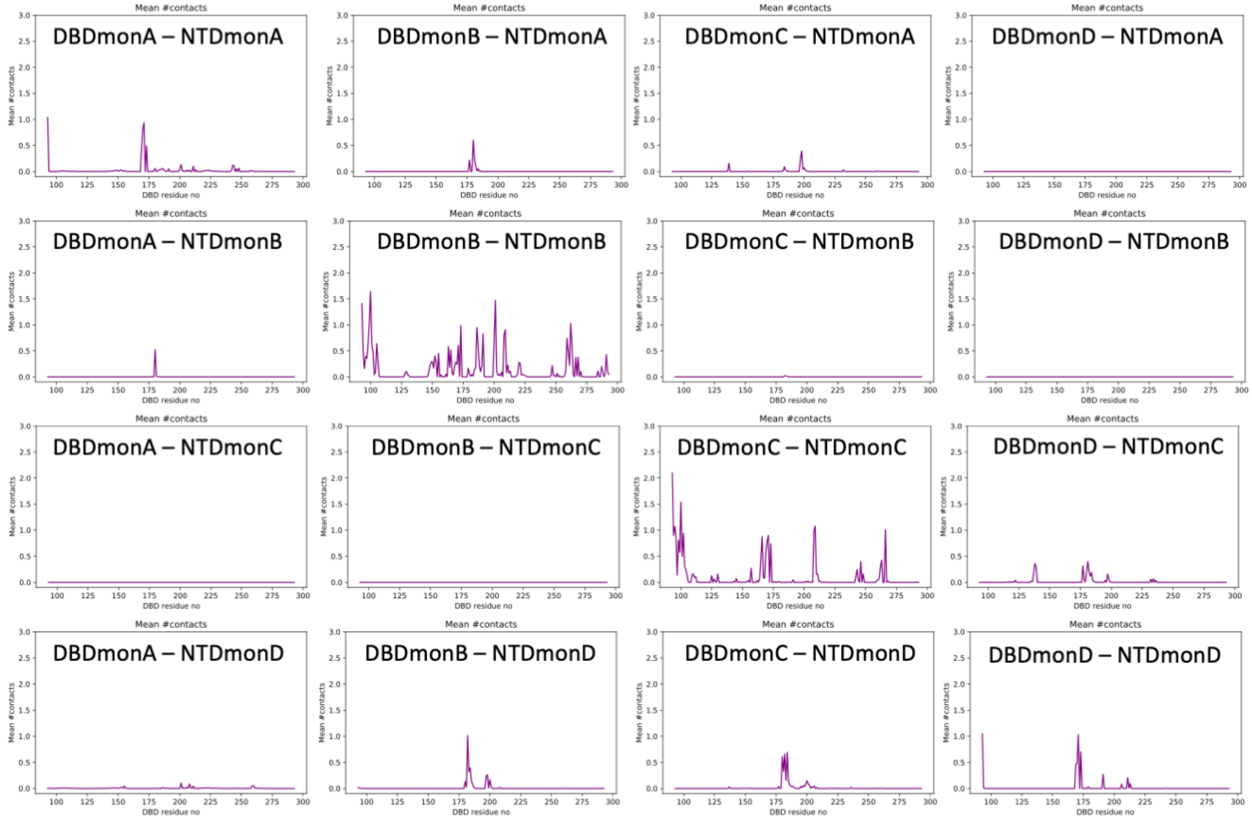
DBDmonA – NTDmonA
DBDmonB – NTDmonA
DBDmonC – NTDmonA
DBDmonD – NTDmonA
DBDmonA – NTDmonB
DBDmonB – NTDmonB
DBDmonC – NTDmonB
DBDmonD – NTDmonB
DBDmonA – NTDmonC
DBDmonB – NTDmonC
DBDmonC – NTDmonC
DBDmonD – NTDmonC
DBDmonA – NTDmonD
DBDmonB – NTDmonD
DBDmonC – NTDmonD
DBDmonD – NTDmonD
Mean #contacts
DBD residue no

Figure S9

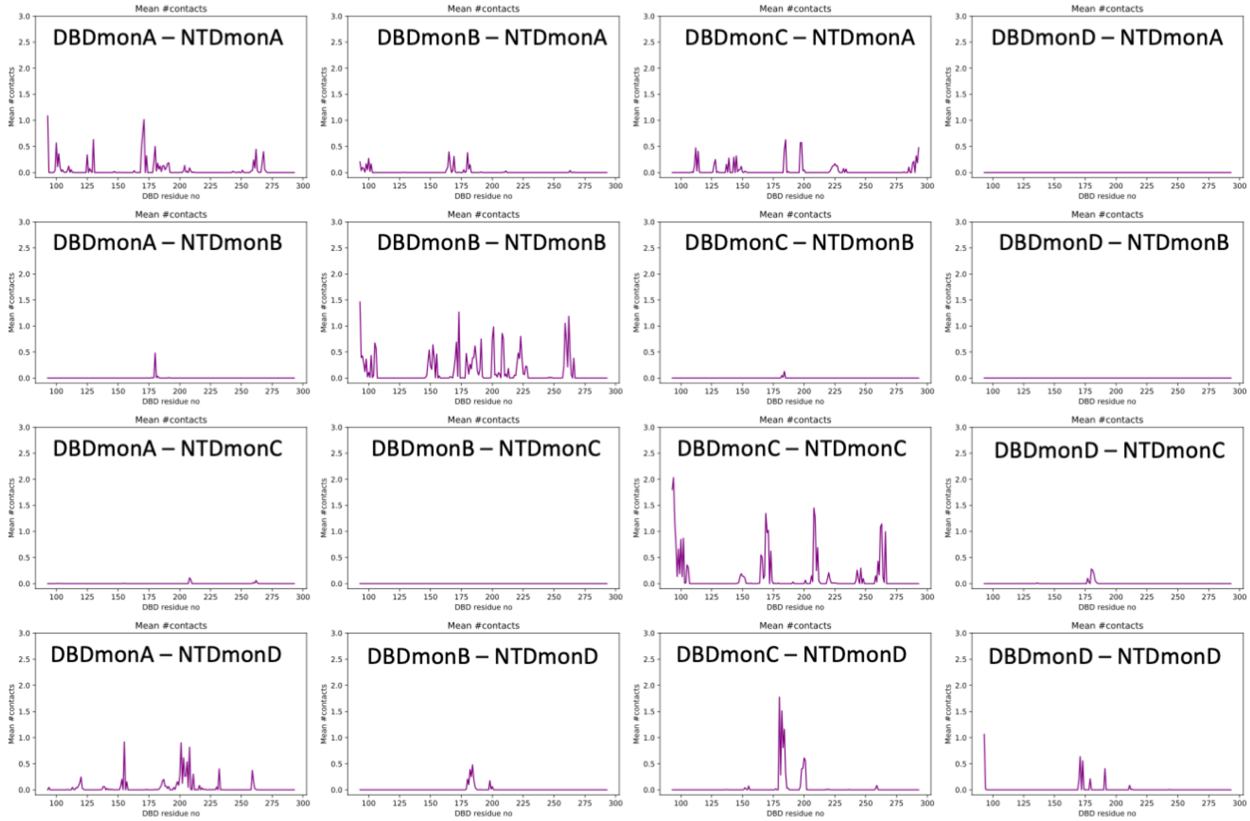
Mean #contacts
DBDmonA – NTDmonA
DBDmonB – NTDmonA
DBDmonC – NTDmonA
DBDmonD – NTDmonA
DBDmonA – NTDmonB
DBDmonB – NTDmonB
DBDmonC – NTDmonB
DBDmonD – NTDmonB
DBDmonA – NTDmonC
DBDmonB – NTDmonC
DBDmonC – NTDmonC
DBDmonD – NTDmonC
DBDmonA – NTDmonD
DBDmonB – NTDmonD
DBDmonC – NTDmonD
DBDmonD – NTDmonD
DBD residue no

Figure S10

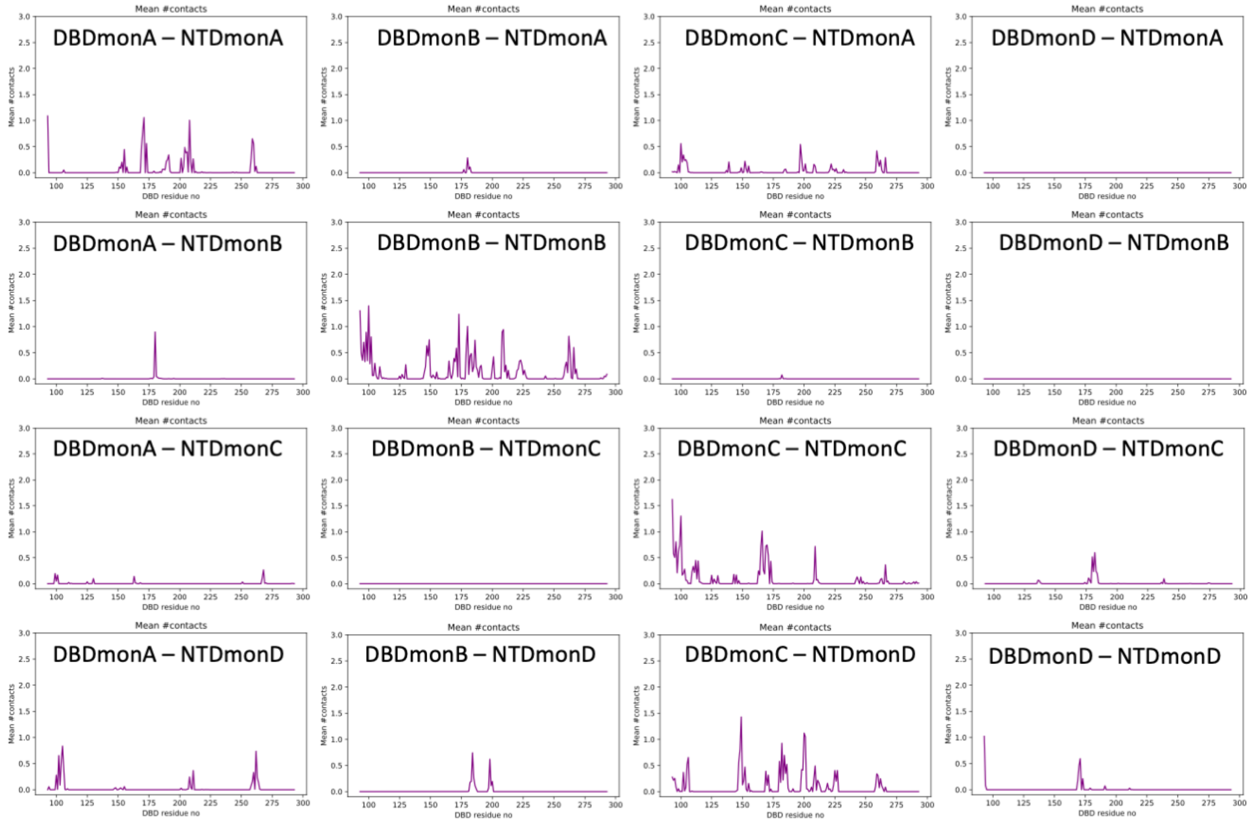
DBDmonA – NTDmonA
DBDmonB – NTDmonA
DBDmonC – NTDmonA
DBDmonD – NTDmonA
DBDmonA – NTDmonB
DBDmonB – NTDmonB
DBDmonC – NTDmonB
DBDmonD – NTDmonB
DBDmonA – NTDmonC
DBDmonB – NTDmonC
DBDmonC – NTDmonC
DBDmonD – NTDmonC
DBDmonA – NTDmonD
DBDmonB – NTDmonD
DBDmonC – NTDmonD
DBDmonD – NTDmonD
Mean #contacts
DBD residue no

Figure S11

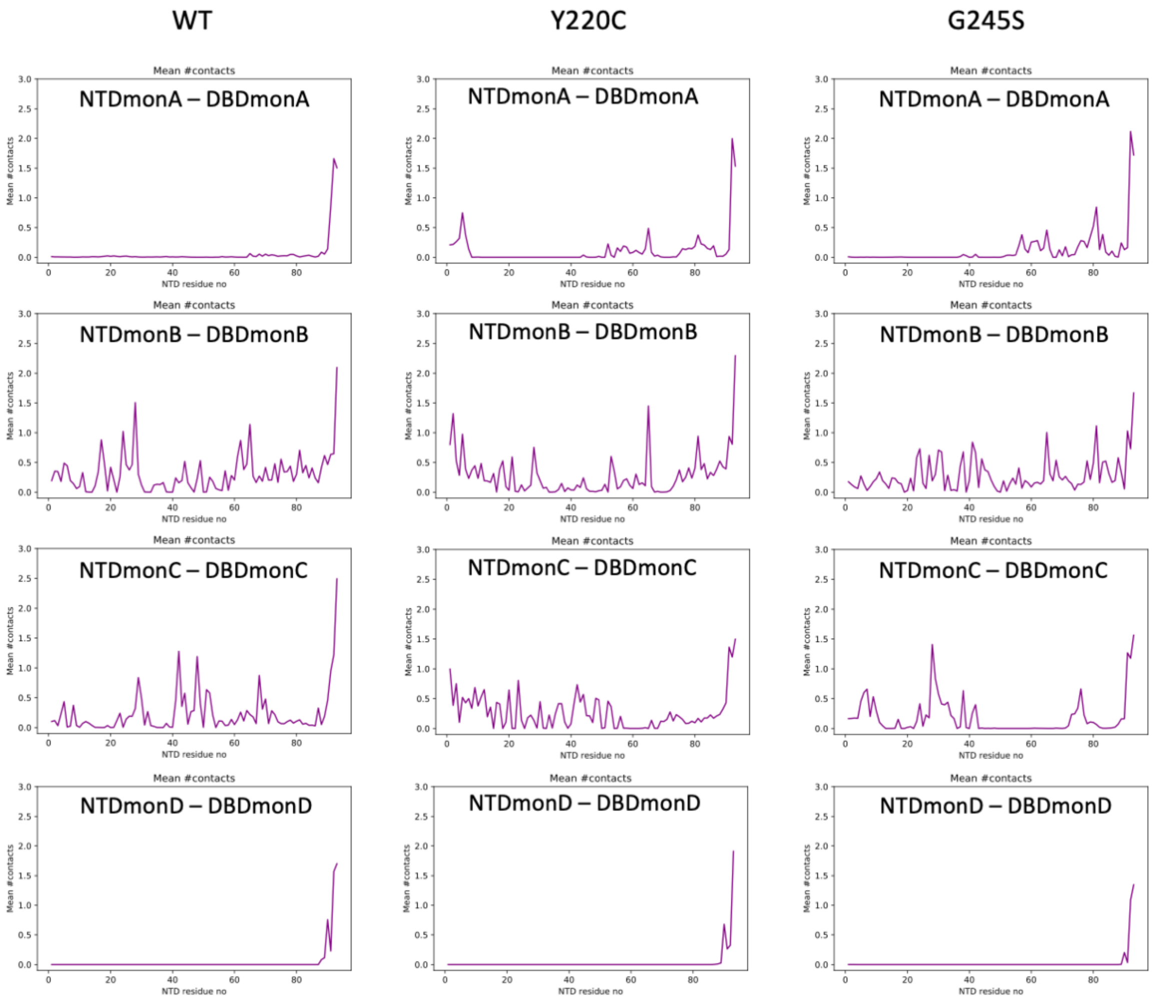
WT
Y220C
G245S
Mean #contacts
NTDmonA – DBDmonA
NTDmonB – DBDmonB
NTDmonC – DBDmonC
NTDmonD – DBDmonD
Mean #contacts
NTD residue no

Figure S12

A

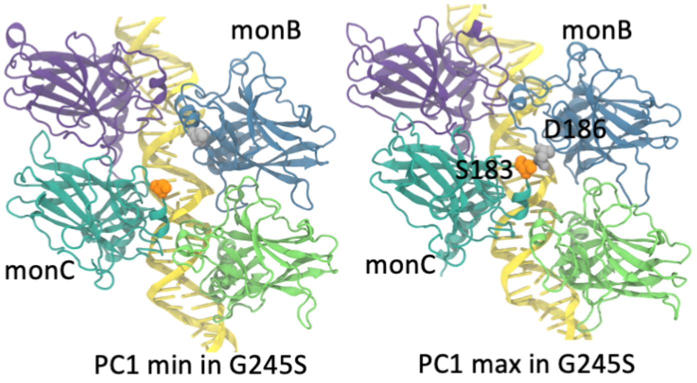


B

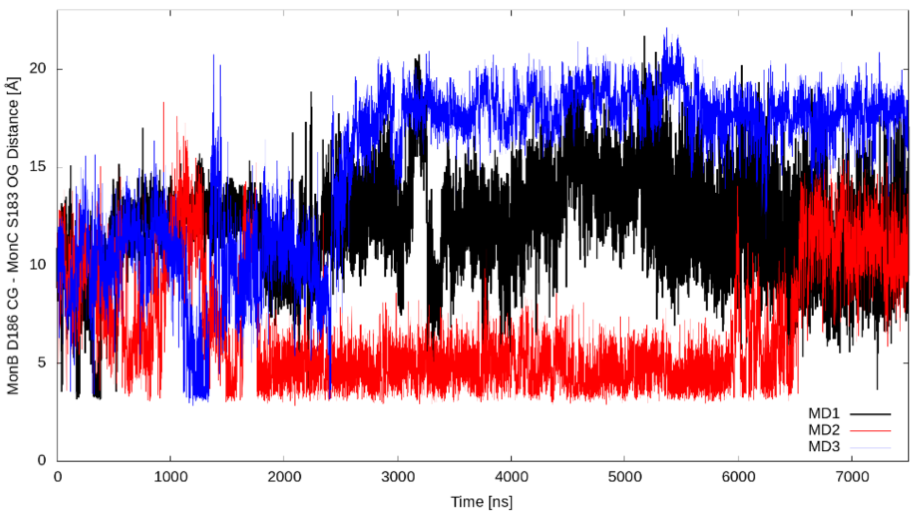

Figure S13

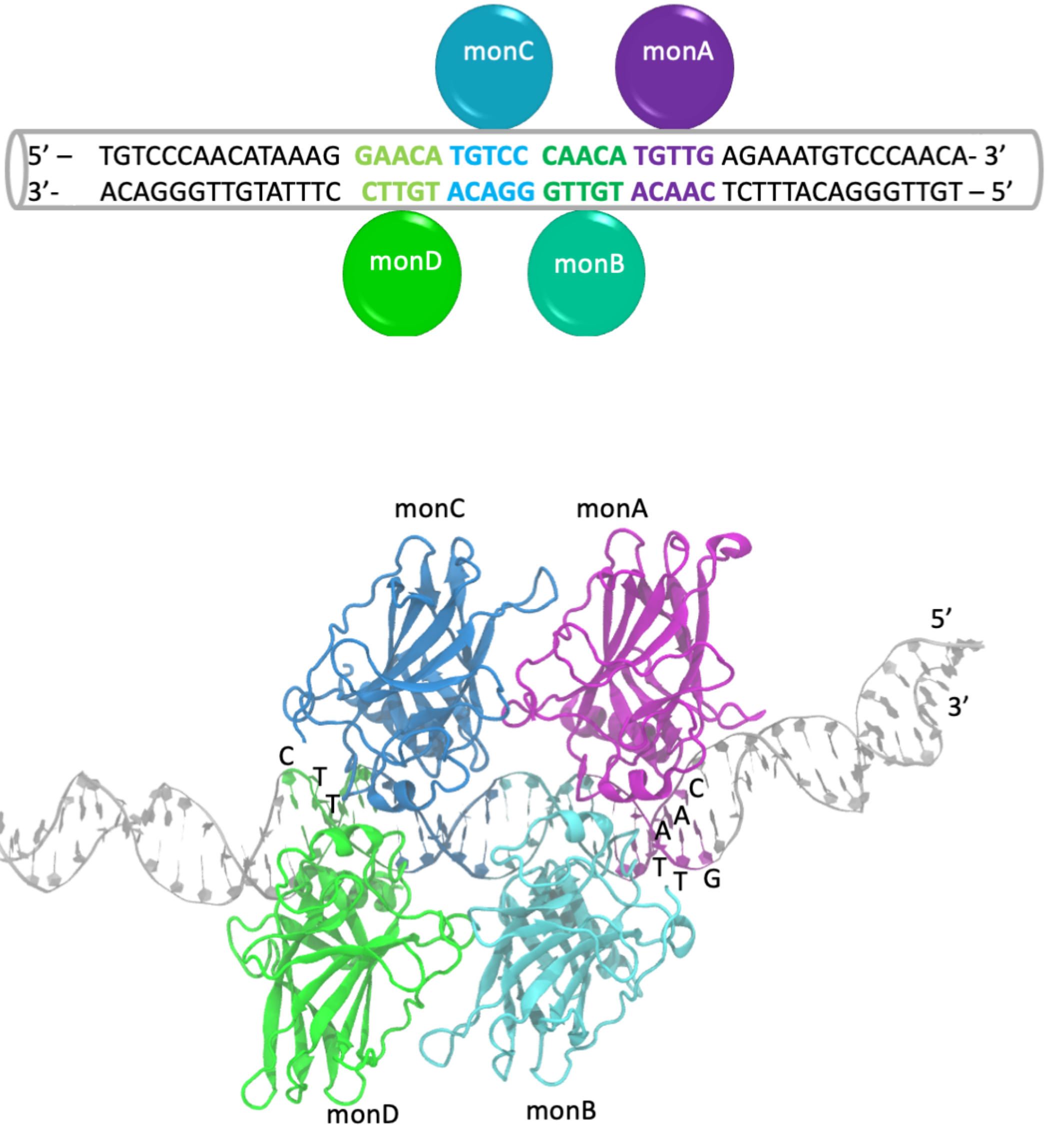
monC
monA
5' – TGTCCCAACATAAAG GAACA TGTCC CAACA TGTTG AGAAATGTCCCAACA- 3'
3'- ACAGGGTTGTATTTC CTTGT ACAGG GTTGT ACAAC TCTTTACAGGGTTGT – 5'
monD
monB
monC
monA
5'
3'
C
T
T
C
A
A
T
T
G
monD
monB

Table S2. The list of WT p53 crystal structures examined & used for comparison in conformational landscapes of Figure 5.

**WT:**

| PDB ID | Chain ID |
|---|---|
| 1GZH | C |
| 1KZY | A,B |
| 1TSR | A,B,C |
| 1TUP | A,B,C |
| 2AC0 | A,B,C,D |
| 2ADY | A,B |
| 2AHI | A,B,D |
| 2ATA | A,B,C,D |
| 2H1L | M,N,O,P,Q,R,S,T,U,V,W,X |
| 2OCJ | A,B,C,D |
| 2XWR | A,B |
| 2YBG | C,D |
| 3KMD | A,B,C,D |
| 3Q05 | A,B,C,D |
| 3TS8 | A,B,C,D |
| 4HJE | A,B,C,D |

Table S3. The list of Y220C p53 crystal structures examined & used for comparison in conformational landscapes of Figure 5.

**Y220C:**

| PDB ID | Chain ID |
|---|---|
| 2J1X | A |
| 2VUK | A,B |
| 2X0U | A |
| 2X0V | A,B |
| 2X0W | A |
| 3ZME | A,B |
| 4AGL | A,B |
| 4AGM | A,B |
| 4AGN | A,B |
| 4AGO | A,B |
| 4AGP | A,B |
| 4AGQ | A,B |
| 5A7B | A,B |
| 5AB9 | A,B |
| 5ABA | A,B |
| 5AOI | A,B |
| 5AOJ | A,B |
| 5AOK | A,B |
| 5AOL | B |
| 5AOM | A,B |
| 5G4M | A,B |
| 5G4N | A,B |
| 5G4O | A,B |
| 6GGA | A,B |
| 6GGB | A,B |
| 6GGC | A,B |
| 6GGD | A,B |
| 6GGE | A,B |
| 6GGF | A,B |

Table S4. The list of G245S p53 crystal structures examined & used for comparison in conformational landscapes of Figure 5.

**G245S:**

| PDB ID | Chain ID |
|---|---|
| 7DHY | A, B, C, D |

Figure S14

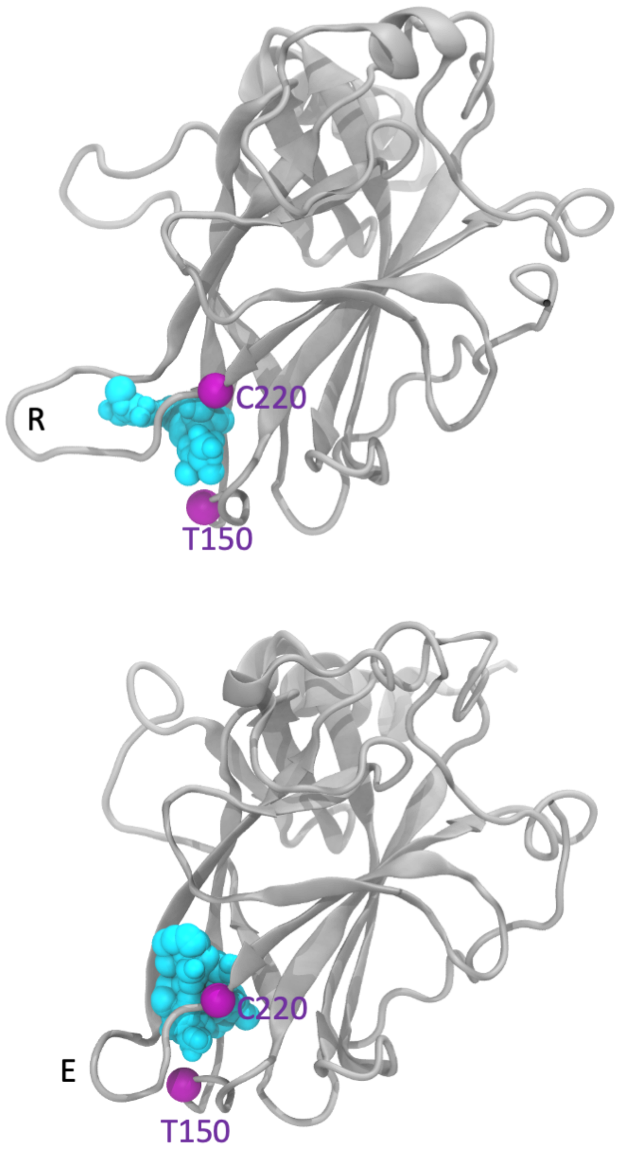

Figure S15

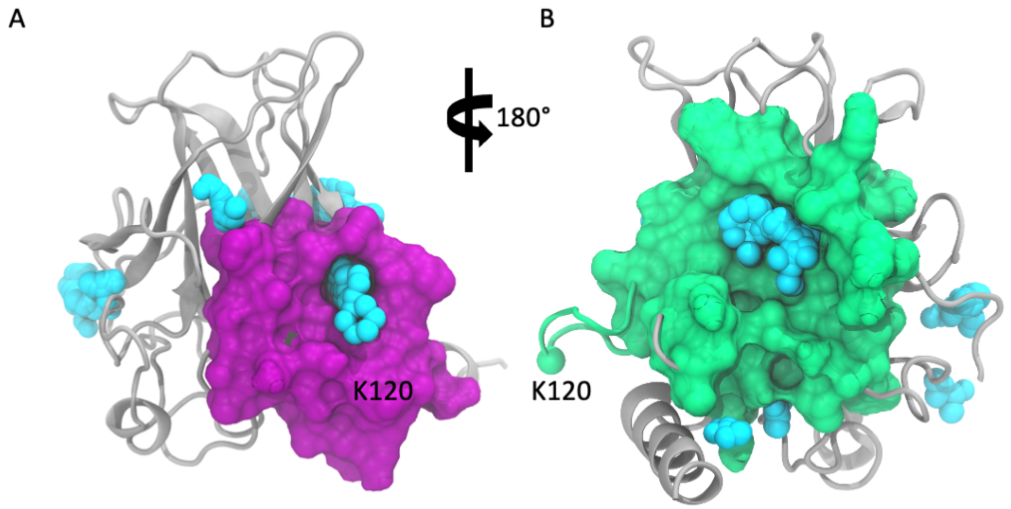

Table S5. Solvent-accessibility of C141 sidechain at L1/S3 pocket (average ∓ standard deviation) and percentage of MD frames with C141 sidechain SASA > 5 Å$^2$

| | **Monomer A** | **Monomer B** | **Monomer C** | **Monomer D** |
|---|---|---|---|---|
| WT C141 SASA (Å$^2$) | 1.16 ∓ 1.60 | 0.80 ∓ 1.19 | 0.84 ∓ 1.50 | 0.54 ∓ 1.00 |
| G245S C141 SASA (Å$^2$) | 0.90 ∓ 1.41 | 0.81 ∓ 1.21 | 0.79 ∓ 1.35 | 0.67 ∓ 1.14 |
| Y220C C141 SASA (Å$^2$) | 0.95 ∓ 1.45 | 0.81 ∓ 1.17 | 0.95 ∓ 1.40 | 0.55 ∓ 1.15 |
| WT C141 exposure % | 3.68 | 1.22 | 2.59 | 0.82 |
| G245S C141 exposure % | 2.52 | 1.31 | 2.09 | 0.99 |
| Y220C C141 exposure % | 2.38 | 1.25 | 2.30 | 0.87 |

Table S6. Solvent-accessibility of C135 sidechain at L1/S3 pocket (average ∓ standard deviation) and percentage of MD frames with C135 sidechain SASA > 5 Å$^2$

| | **Monomer A** | **Monomer B** | **Monomer C** | **Monomer D** |
|---|---|---|---|---|
| WT C135 SASA (Å$^2$) | 0.42 ∓ 1.01 | 0.25 ∓ 0.63 | 0.14 ∓ 0.42 | 0.40 ∓ 0.84 |
| G245S C135 SASA (Å$^2$) | 0.50 ∓ 1.15 | 0.47 ∓ 1.03 | 0.15 ∓ 0.41 | 0.42 ∓ 0.89 |
| Y220C C135 SASA (Å$^2$) | 0.34 ∓ 0.79 | 0.11 ∓ 0.32 | 0.17 ∓ 0.46 | 0.38 ∓ 0.88 |
| WT C135 exposure % | 1.01 | 0.20 | 0.07 | 0.35 |
| G245S C135 exposure % | 1.30 | 0.76 | 0.04 | 0.53 |
| Y220C C135 exposure % | 0.42 | 0.02 | 0.03 | 0.53 |

Table S7. Solvent-accessibility of C182, C229, C275 and C277 sidechain (average ∓ standard deviation) in WT fl-p53 tetramer and percentage of MD frames with cysteine sidechain SASA > 5 Å$^2$

| | **Monomer A** | **Monomer B** | **Monomer C** | **Monomer D** |
|---|---|---|---|---|
| WT C182 SASA (Å$^2$) | 45.71 ∓ 12.60 | 30.58 ∓ 13.98 | 33.87 ∓ 16.33 | 25.63 ∓ 22.37 |
| WT C229 SASA (Å$^2$) | 22.44 ∓ 16.77 | 11.37 ∓ 10.90 | 14.54 ∓ 9.79 | 19.96 ∓ 15.85 |
| WT C275 SASA (Å$^2$) | 1.03 ∓ 2.23 | 0.12 ∓ 0.43 | 0.33 ∓ 0.81 | 0.28 ∓ 1.24 |
| WT C277 SASA (Å$^2$) | 42.64 ∓ 13.01 | 22.23 ∓ 13.70 | 10.51 ∓ 6.02 | 49.16 ∓ 13.40 |
| WT C182 exposure % | 99.96 | 99.50 | 97.90 | 75.99 |
| WT C229 exposure % | 85.78 | 60.94 | 82.96 | 81.39 |
| WT C275 exposure % | 6.57 | 0.18 | 0.62 | 1.00 |
| WT C277 exposure % | 99.99 | 89.88 | 84.69 | 99.92 |

Figure S16

A

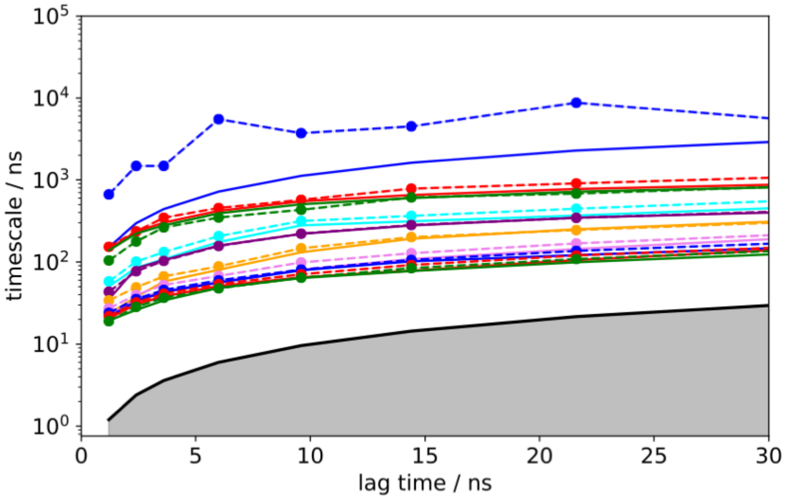


B

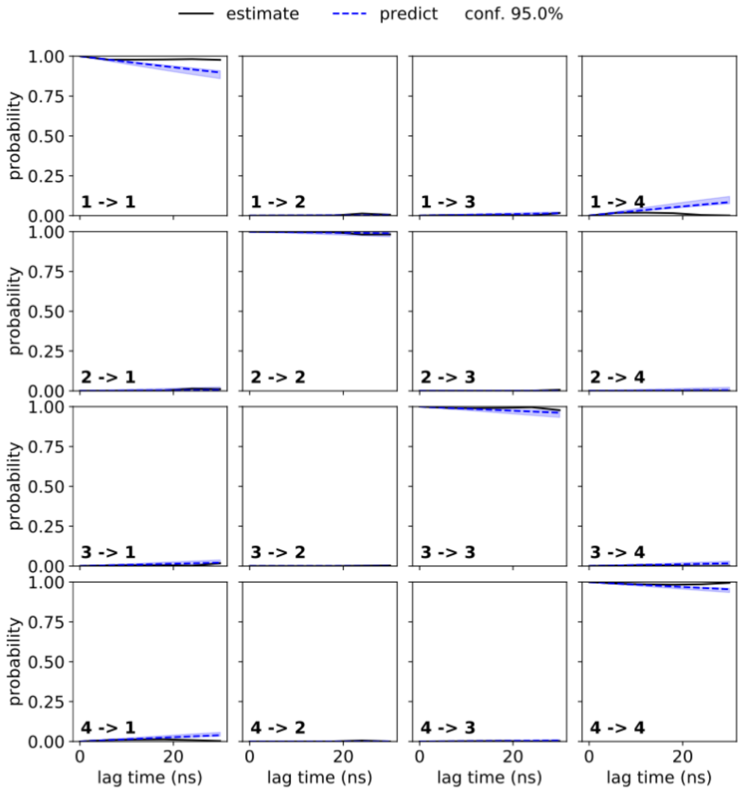

Figure S17

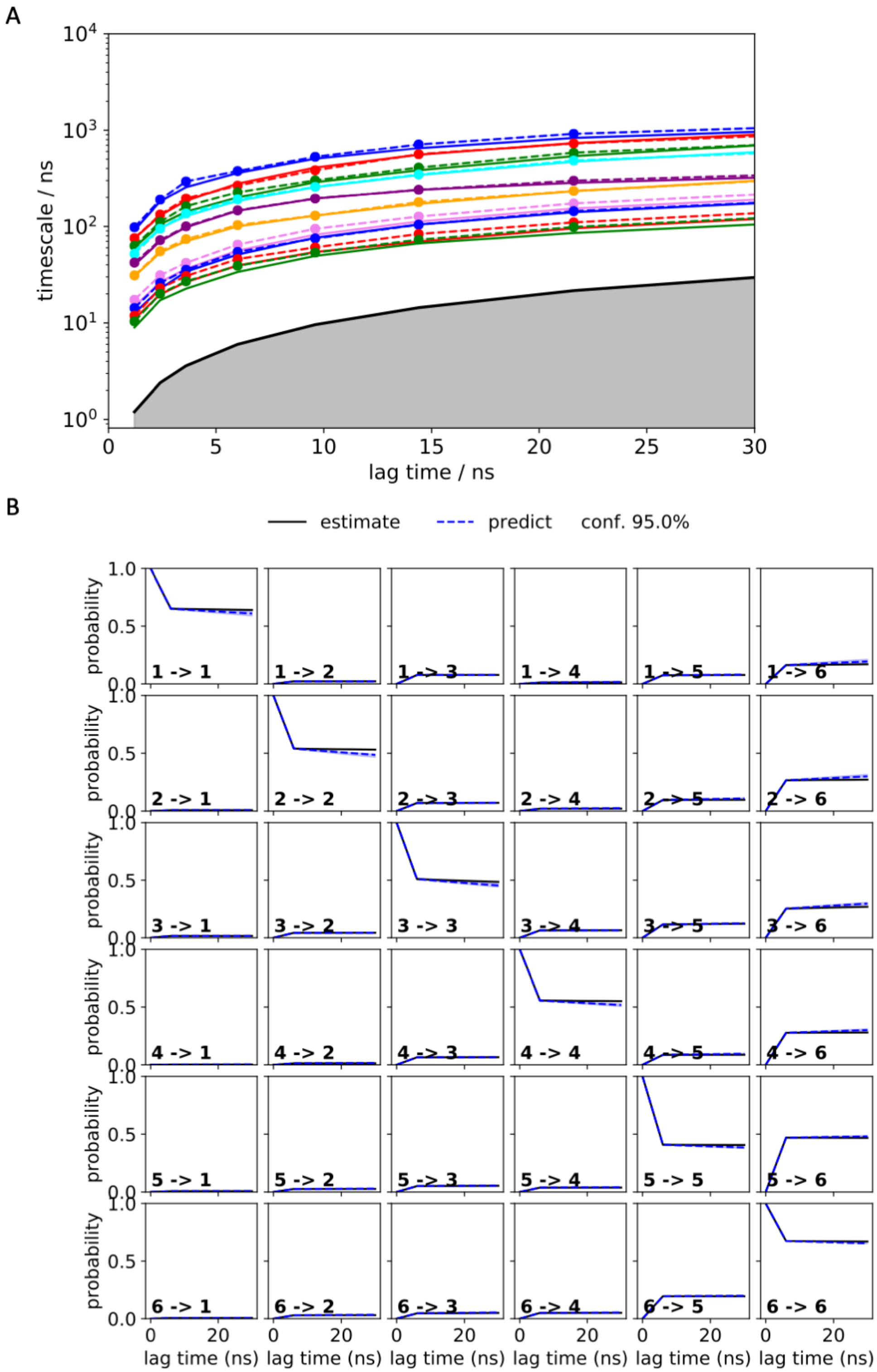
A
timescale / ns
lag time / ns
B
estimate
predict
conf. 95.0%
probability
lag time (ns)
1 -> 1
1 -> 2
1 -> 3
1 -> 4
1 -> 5
1 -> 6
2 -> 1
2 -> 2
2 -> 3
2 -> 4
2 -> 5
2 -> 6
3 -> 1
3 -> 2
3 -> 3
3 -> 4
3 -> 5
3 -> 6
4 -> 1
4 -> 2
4 -> 3
4 -> 4
4 -> 5
4 -> 6
5 -> 1
5 -> 2
5 -> 3
5 -> 4
5 -> 5
5 -> 6
6 -> 1
6 -> 2
6 -> 3
6 -> 4
6 -> 5
6 -> 6

Figure S18

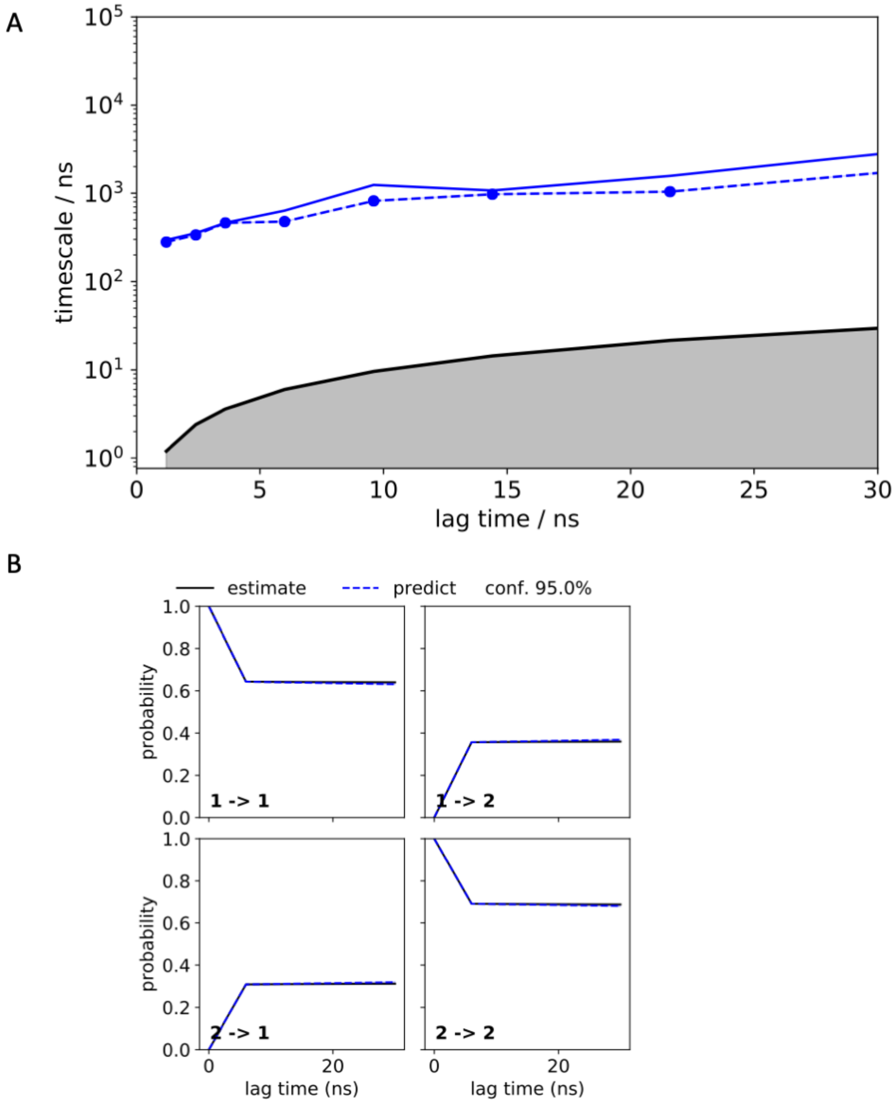

A
$10^5$
$10^4$
$10^3$
$10^2$
$10^1$
$10^0$
timescale / ns
0
5
10
15
20
25
30
lag time / ns
B
estimate
predict
conf. 95.0%
1.0
0.8
0.6
0.4
0.2
0.0
probability
1 -> 1
1 -> 2
2 -> 1
2 -> 2
0
20
lag time (ns)

Figure S19

A

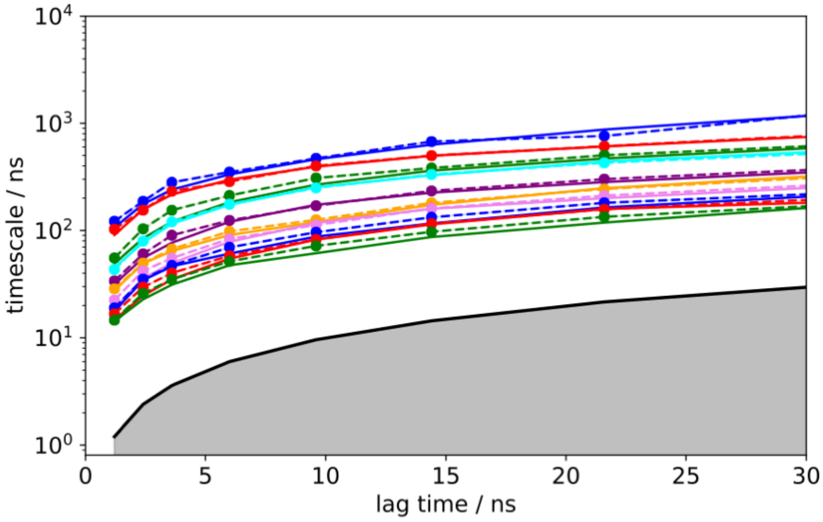


B

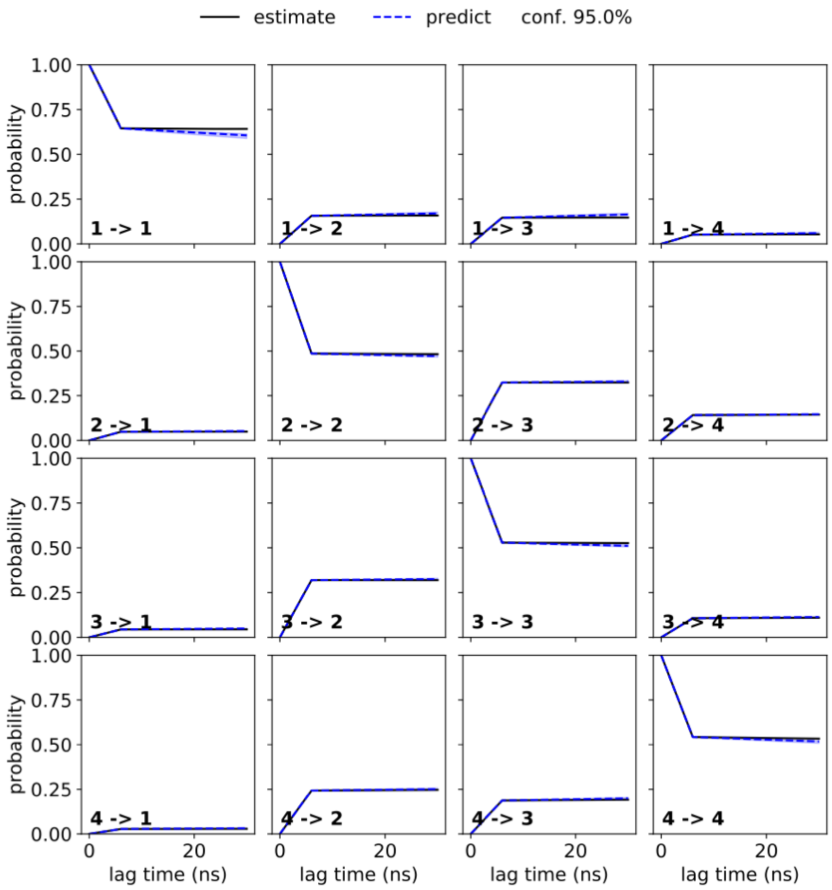

Figure S20

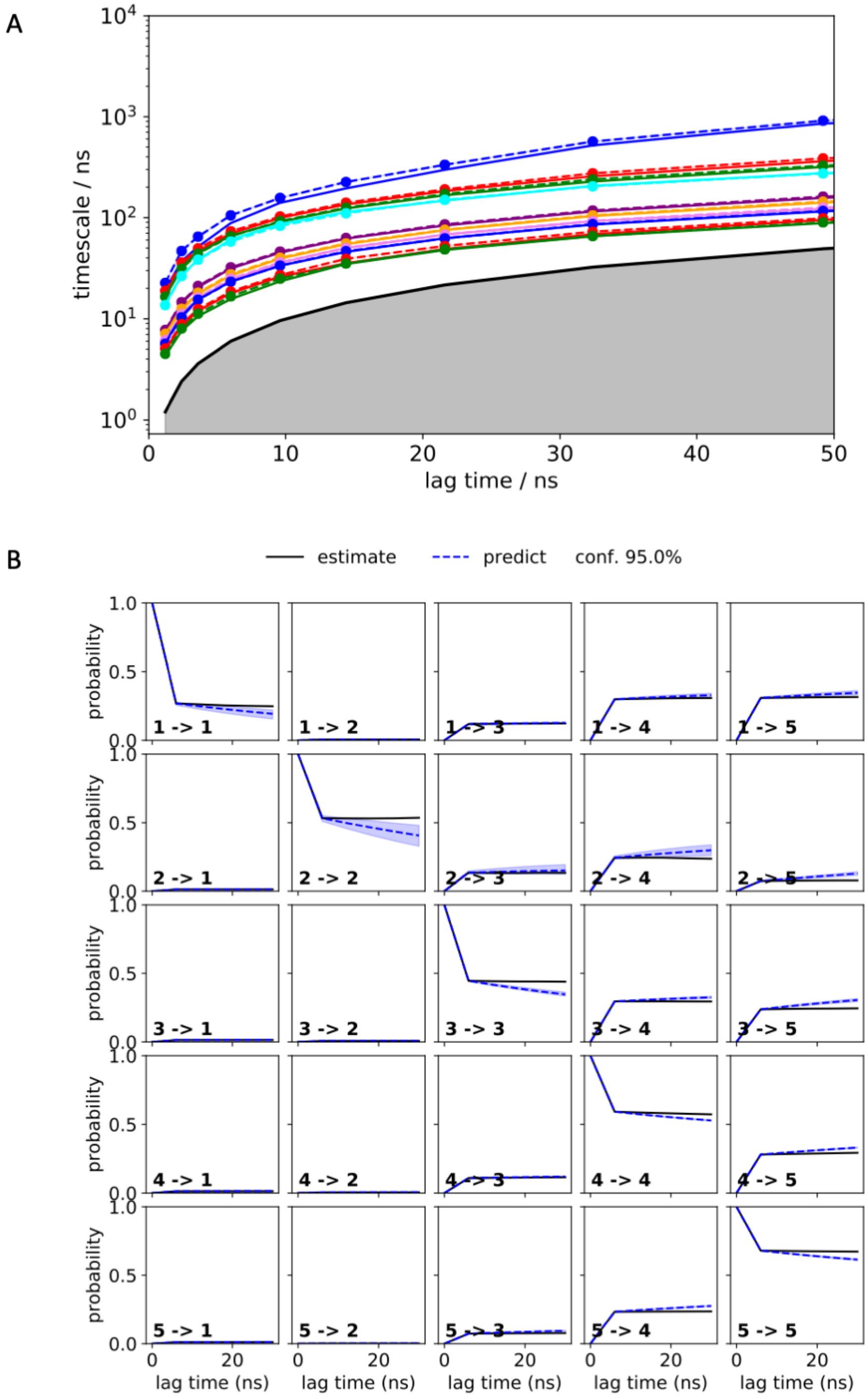
A
timescale / ns
lag time / ns
B
estimate
predict
conf. 95.0%
probability
lag time (ns)
1 -> 1
1 -> 2
1 -> 3
1 -> 4
1 -> 5
2 -> 1
2 -> 2
2 -> 3
2 -> 4
2 -> 5
3 -> 1
3 -> 2
3 -> 3
3 -> 4
3 -> 5
4 -> 1
4 -> 2
4 -> 3
4 -> 4
4 -> 5
5 -> 1
5 -> 2
5 -> 3
5 -> 4
5 -> 5

Figure S21

A

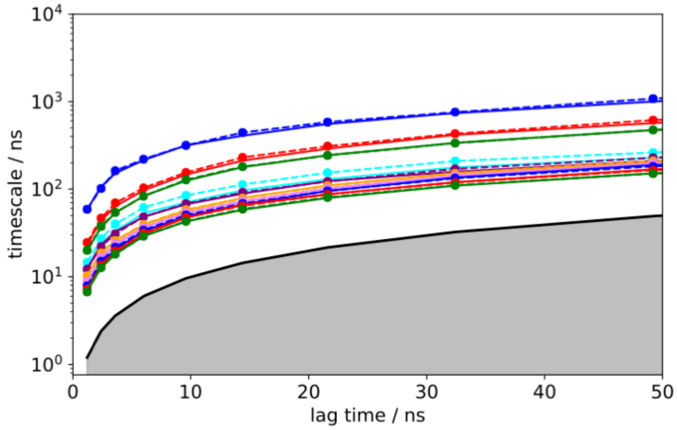


B

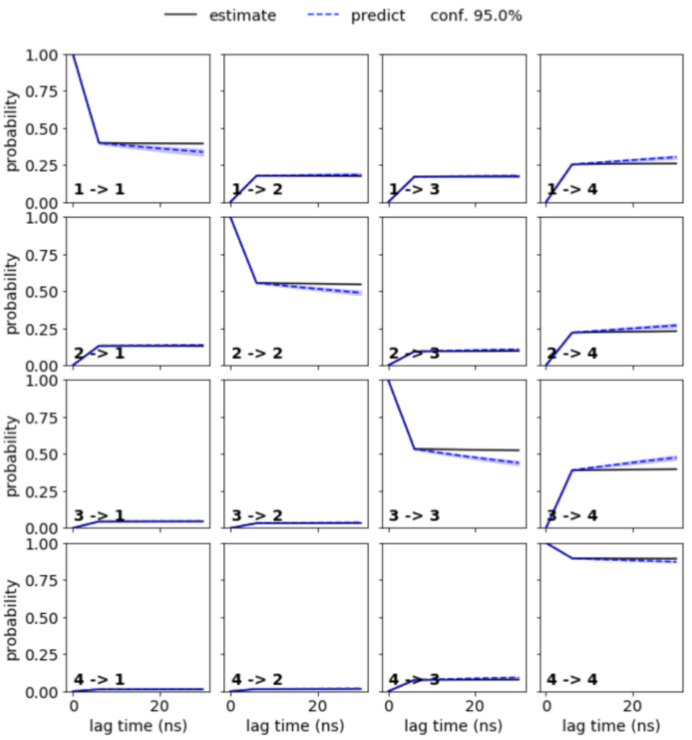

Figure S22

A

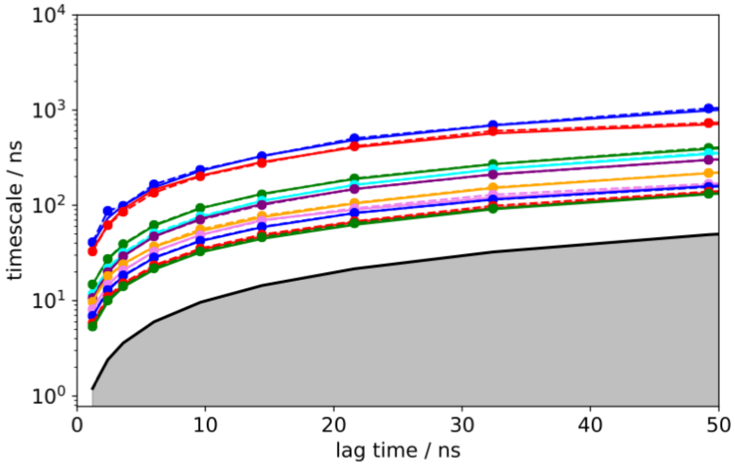


B

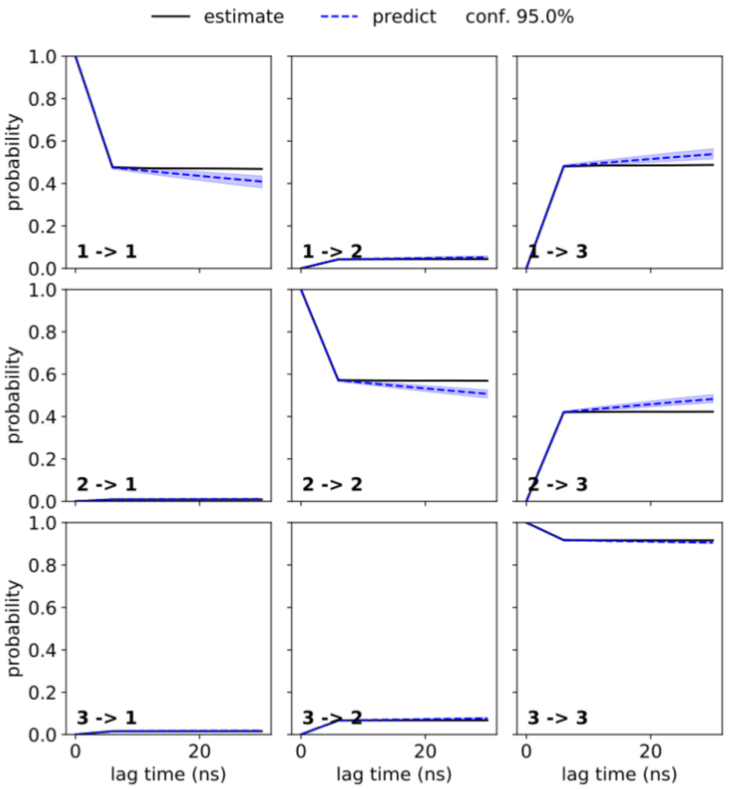

Figure S23

A

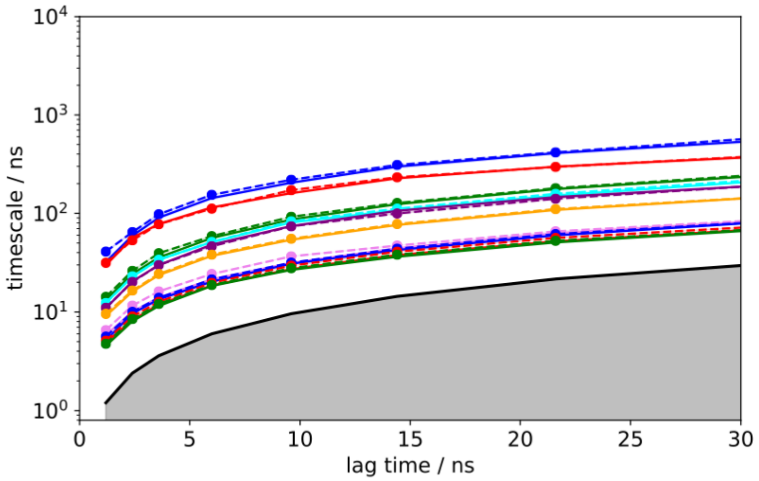


B

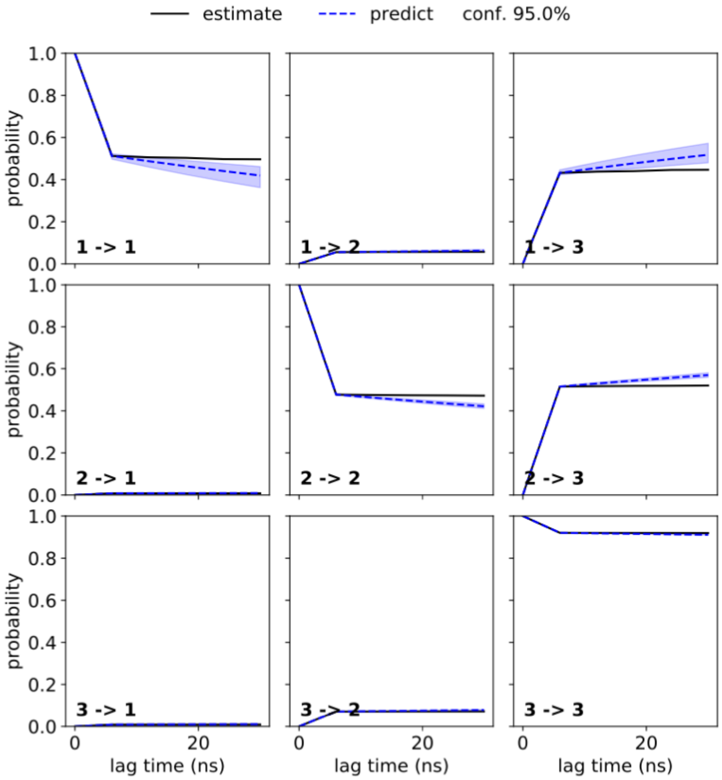
